# Fructose utilization by GM-CSF-differentiated macrophages aggravates autoimmune inflammation *via* MG-derived AGE–RAGE signaling

**DOI:** 10.64898/2026.01.26.701899

**Authors:** Yeon Jun Kang, Sihyun Chae, Jaemoon Koh, Woogil Song, Yufei Li, Jeong-Won Kim, Hee Young Kim, Dongjin Jeong, Jong-Wan Park, Eun Young Lee, Doo Hyun Chung, Seong Wook Kang, Jin Kyun Park, Jeong Seok Lee, Joo-Youn Cho, Won-Woo Lee

**Affiliations:** Laboratory of Autoimmunity and Inflammation, Department of Biomedical Sciences, Seoul National University College of Medicine, Seoul 03080, Republic of Korea; Department of Microbiology and Immunology, Seoul National University College of Medicine, Seoul 03080, Republic of Korea; Department of Clinical Pharmacology and Therapeutics, Seoul National University College of Medicine and Hospital, Seoul 03080, Republic of Korea; Department of Biomedical Sciences, Seoul National University College of Medicine, Seoul 03080, Republic of Korea; Department of Pathology, Seoul National University College of Medicine, Seoul 03080, Republic of Korea; Graduate School of Medical Science and Engineering, Korea Advanced Institute of Science and Technology (KAIST), Daejeon, 34141, Republic of Korea; Institute of Endemic Diseases, Seoul National University Medical Research Center. Seoul 03080, Republic of Korea; Laboratory of Immune Regulation, Department of Biomedical Sciences, Seoul National University College of Medicine, Seoul 03080, Republic of Korea; Department of Pharmacology, Seoul National University College of Medicine, Daehak-ro 103, Jongno-gu, Seoul 03080, Republic of Korea; Division of Rheumatology, Department of Internal Medicine, Seoul National University College of Medicine, Seoul 03080, Republic of Korea; Department of Internal Medicine, Chungnam National University College of Medicine, 282 Munhwa-ro, Jung-gu, Daejeon 35015, Republic of Korea; Seoul National University Cancer Research Institute; Ischemic/Hypoxic Disease Institute, Seoul National University Medical Research Center; Seoul National University Hospital Biomedical Research Institute, Seoul 03080, Republic of Korea

**Keywords:** fructose metabolism, ketohexokinase (KHK), MG-AGE, RAGE signaling, GM-CSF macrophage

## Abstract

Excessive fructose intake is increasingly associated with metabolic and inflammatory pathologies; however, the direct impact of fructose metabolism on immune cell function remains insufficiently understood. Here, we demonstrate that GM-CSF–differentiated macrophages markedly upregulate the fructose-specific transporter GLUT5, enabling efficient fructose utilization through two complementary metabolic pathways. Using ^13^C-fructose tracing, we identified distinct carbon fluxes *via* a hexokinase (HK)-dependent glycolytic route and a ketohexokinase (KHK)-ALDOB–mediated fructolytic route. The HK-dependent pathway sustains glycolytic activity under glucose-limiting conditions, stabilizing HIF-1α and preserving pro-inflammatory gene expression. Conversely, the KHK–ALDOB axis increases dihydroxyacetone phosphate (DHAP) production, leading to methylglyoxal-derived advanced glycation end-products (MG-AGEs) that engage receptor for AGE (RAGE) signaling. This activation enhances MMP9 expression in macrophages and drives Th17 differentiation in CD4⁺ T cells, amplifying inflammatory and tissue-remodeling processes. *In vivo,* fructose intake exacerbates autoimmune arthritis in SKG mice, and elevated fructose-driven glycolysis and MG-AGE accumulation are observed in monocytes from patients with rheumatoid arthritis. These results link fructose-driven carbonyl stress to macrophage effector programs and downstream T cell polarization. Collectively, these findings uncover a dual fructose metabolic program that integrates nutrient stress with persistent inflammation via the MG-AGE–RAGE axis and underscore the immunometabolic risks of excessive fructose exposure.

## INTRODUCTION

In recent decades, there has been a significant rise in the prevalence of non-communicable diseases (NCDs), such as cardiovascular diseases, type 2 diabetes, metabolic disorders, non-alcoholic fatty liver disease, and autoimmune diseases^1, 2, 3, 4^. These diseases are largely linked to chronic inflammation and lifestyle factors, particularly dietary habits^5^. A prominent feature of the Western diet is excessive consumption of dietary fructose, largely due to increased intake of sucrose (a major component of refined sugar) and high fructose corn syrup (HFCS), commonly found in processed foods and sweetened beverages^6, 7, 8^. Fructose’s unique metabolic properties may pose particular health risks, with high fructose diets being considered key contributors to the global epidemics of metabolic syndrome and chronic systemic inflammation^9^. Notably, high fructose consumption has been associated with chronic inflammatory conditions such as rheumatoid arthritis (RA)^10^, although the exact mechanisms linking fructose to these diseases remain unclear.

The majority of dietary fructose is absorbed passively *via* the fructose-specific transporter SLC2A5 in the intestine and is primarily metabolized in the liver, where it is converted into glucose, lactate, and glycogen^11, 12^. Under normal conditions, fructose levels in peripheral plasma are around 0.04 mM, but these levels increase rapidly by up to 10-fold after fructose intake, returning to baseline within a few hours^9^. Elevated fructose levels are observed in certain pathological conditions, particularly in patients with acute myeloid leukemia, where fructose concentrations can reach as high as 5 mM in the bone marrow^13^. Emerging evidence suggests that several cancer cells adapt their metabolism to utilize fructose as an alternative energy source when glucose is limited in the tumor microenvironment^14, 15, 16, 17^. This fructose utilization is further enhanced by the upregulation of molecules and enzymes involved in fructose uptake and metabolism^9, 18, 19^. Fructose is metabolized through glycolysis, either by ketohexokinase (KHK), which produces fructose-1-phosphate (a substrate for aldolase B), or by hexokinase (HK), which converts it to fructose-6-phosphate, though this process is less efficient than glucose metabolism^20, 21, 22, 23^.

Immune responses necessitate considerable metabolic reprogramming, particularly through enhanced glycolysis, to fulfill the substantial energy requirements of activated immune cells. While glucose is typically the primary energy source for immune cells, recent studies have shown that monocytes and macrophages demonstrate metabolic plasticity and can utilize various energy sources. However, the impact of elevated fructose, the second most abundant dietary sugar in humans, on the immune system has not been extensively studied. Recent research has indicated that fructose promotes glutaminolysis in monocytes, enhancing IL-1β production in an acute inflammatory mouse model^24^. Additionally, in macrophages, fructose has been shown to regulate cytosolic Ca²⁺ levels, influencing inflammatory responses in immune-suppressive tumor microenvironments^25^. These findings suggest that fructose may play a broader role in modulating macrophage metabolism, though the specific metabolic pathways involved remain unclear.

In this study, we explore the role of fructose in metabolic pathways and its effects on immune responses in inflammatory macrophages, specifically those differentiated with GM-CSF. Our results show that SLC2A5 (GLUT5), a fructose-specific transporter, is upregulated in GM-CSF macrophages facilitating fructose uptake. Fructose acts as an alternative carbohydrate source to support hexokinase (HK)-dependent glycolytic flux, while also undergoing fructolysis *via* ketohexokinase (KHK) and aldolase B (ALDOB), both of which are overexpressed in GM-CSF macrophages. This metabolic reprogramming leads to the production of methylglyoxal-derived advanced glycation end-products (MG-AGEs), which enhance inflammatory responses through receptor for AGEs (RAGE) signaling in both macrophages and T cells. *In vivo*, fructose supplementation exacerbates disease severity in the autoimmune arthritis mouse model, underscoring its pathophysiological relevance. Overall, our findings suggest that fructose metabolism plays a critical role in driving immune cell-mediated inflammation and may serve as a potential therapeutic target for chronic inflammatory diseases like rheumatoid arthritis.

## RESULTS

### The expression of SLC2A5, a fructose-specific transporter, is markedly increased in inflammatory macrophages

In patients with rheumatoid arthritis (RA), synovial monocytes and macrophages exhibit pronounced activation, potentially reflecting an adaptive regulation of metabolic homeostasis to sustain the elevated energetic demands of these immune cells^26, 27^. Glucose serves as the primary carbon source for cellular metabolism; however, reanalysis of our previous microarray dataset (E-MTAB-6187) revealed that among sugar transporters, SLC2A5 expression was significantly upregulated in synovial monocytes from RA patients compared with peripheral blood monocytes from the same individuals (Fig. 1A). This finding was further validated in an independent cohort, where purified CD14⁺ monocytes from RA synovial fluid displayed higher SLC2A5 expression than their peripheral counterparts (Fig. 1B). Thus, this suggests that the synovial microenvironment in RA plays a pivotal role in the transcriptional activation of SLC2A5. Moreover, treatment of healthy donor-derived monocytes with 10% RA synovial fluid induced both mRNA and protein expression of SLC2A5 (GLUT5) (Fig. 1C, D), confirming that soluble mediators in the RA synovium can upregulate this transporter.

**Figure 1.**
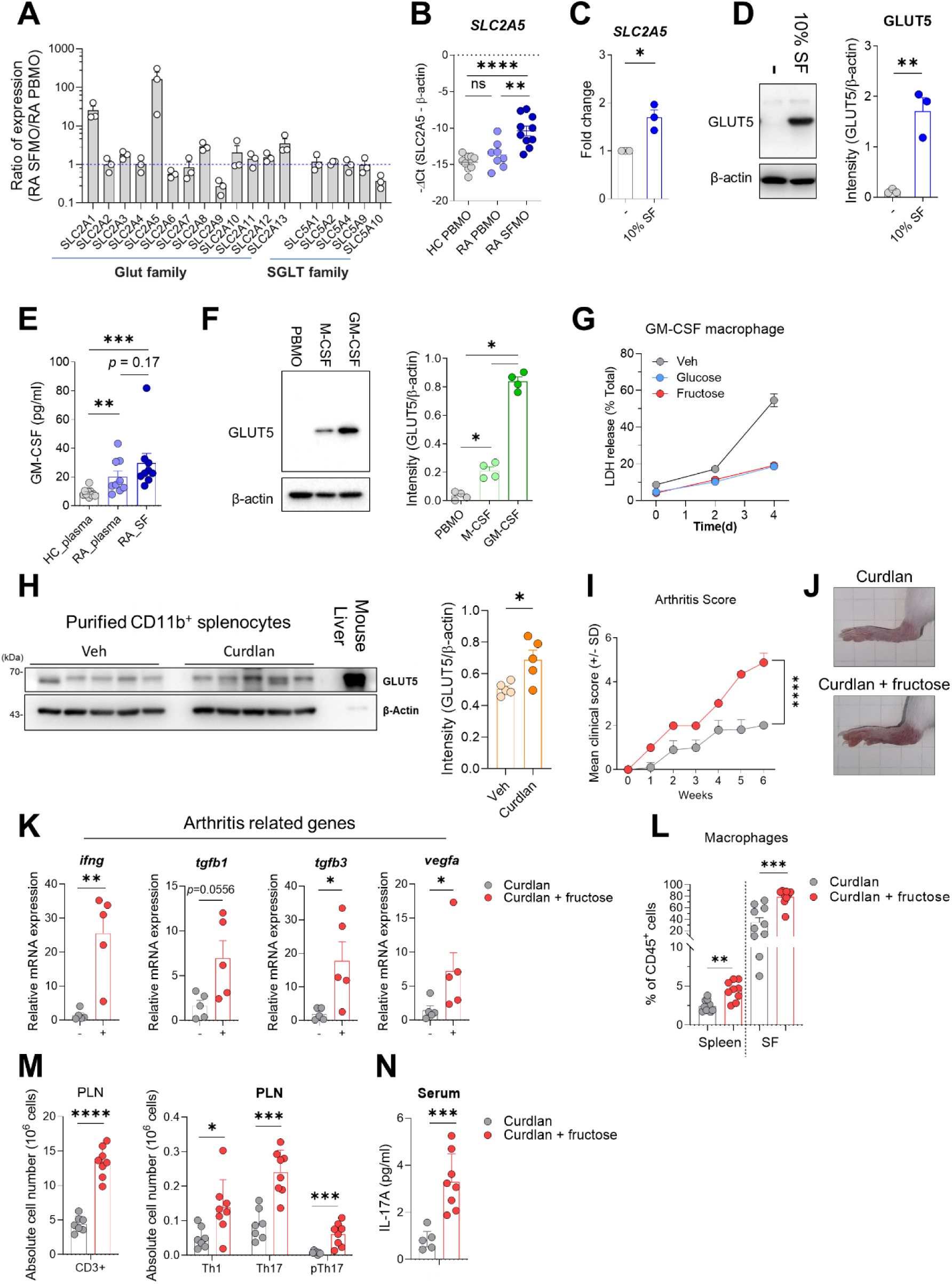
GM-CSF promotes fructose metabolism in human monocytes. **A.** Microarray analysis of monocytes isolated from peripheral blood mononuclear cells (PBMCs) and synovial fluid mononuclear cells (SFMCs) of rheumatoid arthritis (RA) patients (n=3). **B.** Quantitative RT-PCR analysis of *SLC2A5* mRNA in monocytes derived from healthy control PBMCs (HC PBMO), RA PBMCs (RA PBMO), and RA synovial fluid (RA SFMO). **C-D.** *SLC2A5* expression in healthy monocytes cultured for 24 h with or without RA synovial fluid, analyzed by RT-qPCR (C) and immunoblotting (D). Representative blots are shown (D, left). **E.** GM-CSF concentrations measured in plasma from age- and gender-matched healthy controls, RA patient plasma, and RA synovial fluid. **F**. Immunoblot analysis of GLUT5 protein expression in monocytes differentiated with M-CSF (50 ng/ml) or GM-CSF (50 ng/ml) for six days; representative blot shown (left). **G.** Cell viability of GM-CSF macrophages cultured with or without 10 mM fructose or glucose, assessed by LDH release over time (n=5). **H–L**. SKG mice were administered curdlan to induce arthritis and received fructose in drinking water starting three days before induction and continued until sacrifice. (H) Immunoblot analysis of GLUT5 in splenic CD11b⁺ cells at six weeks post-induction (n=5 per group); mouse liver served as a positive control. (I) Clinical arthritis scores (n=5 per group). (J) Representative images of ankle joints. (K) RT-qPCR analysis of synovial mRNA expression of arthritis-related genes. (L) Frequency of CD11b⁺F4/80⁺ macrophages among CD45⁺ cells in spleen and synovial fluid (n=10 per group). **M.** Absolute numbers of CD3⁺ T cells, Th1 (IFN-γ⁺), Th17 (IL-17⁺), and pathogenic Th17 (IFN-γ⁺IL-17⁺) cells in popliteal lymph nodes. **N.** Serum IL-17 levels quantified by ELISA. Band intensities were normalized to β-actin. Data are presented as mean ± SEM. \**p*<0.05, \*\**p*<0.01, \*\*\**p*<0.001, \*\*\*\**p*<0.0001 (C, D: unpaired Student’s t-test; E, F, K, L: Mann–Whitney U test; I: two-way ANOVA).

Synovial monocytes migrating from the circulation serve as precursors for inflammatory macrophages and osteoclasts within RA joints^28^. M-CSF and GM-CSF are critical for macrophage differentiation, with GM-CSF driving proinflammatory or pathogenic phenotypes and M-CSF primarily supporting homeostatic functions^29^. In line with this, GM-CSF levels were substantially higher in the synovial fluid of RA patients compared with plasma from both RA patients and healthy controls (Fig. 1E). We next examined SLC2A5 expression in monocytes and in macrophages differentiated under M-CSF or GM-CSF conditions (hereafter referred to as M-CSF macrophages and GM-CSF macrophages, respectively). While resting monocytes showed negligible SLC2A5 expression, GM-CSF macrophages displayed pronounced upregulation of SLC2A5 (Fig. 1F and Supplementary Fig. 1A) accompanied by increased expression of CXCL10, a canonical GM-CSF macrophage marker (Supplementary Fig. 1B). Various SLC transporters are transcriptionally regulated by c-Myc to support cellular metabolic demands^30, 31^, and the SLC2A5 promoter was found to contain putative MYC response elements. Consistently, pharmacological inhibition of c-Myc with 10058-F4 markedly attenuated GLUT5 induction in GM-CSF macrophages (Supplementary Fig. 1C).

To determine whether fructose could serve as an alternative energy substrate to glucose in macrophages, we compared the viability of GM-CSF macrophages cultured with fructose or glucose supplementation^32^. In glucose-deprived conditions, lactate dehydrogenase (LDH) release was significantly elevated on day 4, indicating reduced cell viability. Supplementation with fructose, however, restored macrophage viability to a level comparable to that observed with glucose (Fig. 1G). These findings suggest that fructose, imported through SLC2A5, can act as an alternative metabolic substrate supporting the survival of inflammatory GM-CSF macrophages.

To further investigate the pathogenic role of fructose metabolism in RA, we administered fructose *via* drinking water to SKG mice, a murine model of autoimmune arthritis. Similar to human macrophages, bone marrow-derived macrophages (BMDMs) from mice exhibited elevated GLUT5 expression following GM-CSF differentiation (Supplementary Fig. 1D). Autoimmune responses in SKG mice elicited through intraperitoneal curdlan injection induced polarization of CD11b⁺ cells into F4/80⁺ macrophages (Supplementary Fig. 2A), and GLUT5 expression was upregulated in splenic CD11b⁺ cells of curdlan-treated mice (Fig. 1H). Mice receiving 15% (w/v) fructose, an intake comparable to that in commercially available soft drinks, developed more severe arthritis, accompanied by enhanced expression of arthritis-associated genes in synovial immune cells (Fig. 1I–K and Supplementary Fig. 2B-C)^33^. Fructose administration also increased the frequency of macrophages in synovial fluid and, to a lesser extent, in the spleen (Fig. 1L). In addition to macrophage activation, CD4⁺ T cells exhibited heightened inflammatory responses, characterized by increased frequencies of IL-17⁺ and/or IFN-γ⁺ cells and elevated serum IL-17A concentrations (Fig. 1M, N). Importantly, the fructose concentrations and treatment durations used did not induce hepatic abnormalities such as steatosis, ballooning, or fibrosis (Supplementary Fig. 2D-F), indicating that the exacerbated arthritis observed was not secondary to fructose-induced liver inflammation. Collectively, these findings demonstrate that GLUT5 is selectively upregulated in GM-CSF–dependent inflammatory macrophages and that GLUT5-mediated fructose uptake contributes to the progression of autoimmune arthritis by enhancing inflammatory cell metabolism within the synovial microenvironment.

### Fructose possesses the capacity to sustain inflammatory activation in GM-CSF macrophages by functionally substituting for glucose

To determine whether fructose can replace glucose in supporting macrophage inflammatory responses, we assessed cytokine production in LPS-stimulated GM-CSF macrophages cultured with either fructose or glucose as the carbohydrate source. Fructose treatment increased IL-1β secretion relative to carbohydrate-free conditions, reaching levels comparable to those observed with glucose supplementation. Similarly, IL-6 production was enhanced in fructose-treated macrophages, albeit to a lesser extent than with glucose. In contrast, IL-23 production remained unaffected by treatment with either sugar (Fig. 2A).

**Figure 2.**
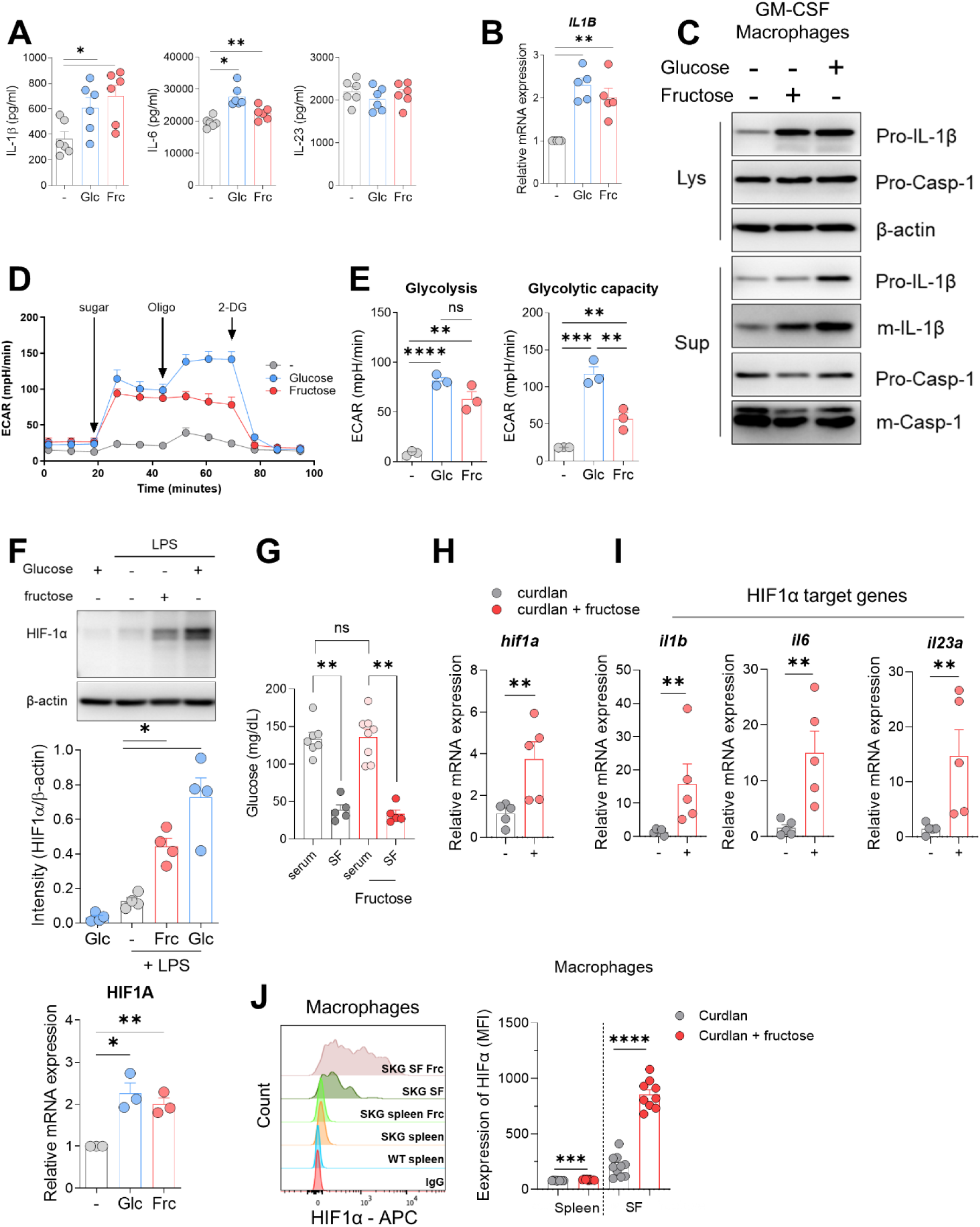
Fructose enhances glycolysis and pro-inflammatory cytokine production in human macrophages. **A.** GM-CSF human macrophages were stimulated with LPS (50 ng/mL) for 18 h in glucose-, fructose- (10 mM), or sugar-free media. IL-1β, IL-6, and IL-23 levels were quantified by ELISA. For IL-1β detection, 4 mM ATP was added during the final 2 h (n = 6). **B.** IL1B mRNA expression was analyzed by RT-qPCR under identical conditions (n = 5). **C.** Immunoblotting of cell lysates and supernatants from (A) was performed to assess pro- and mature (m) forms of IL-1β and caspase-1 (Casp-1). Band intensities were quantified by densitometry and normalized to β-actin (n = 5). **D–E.** Glycolytic activity was assessed by Seahorse extracellular flux analysis of ECAR following sequential injection of glucose, fructose (10 mM), or no sugar, followed by oligomycin (2 µM) and 2-DG (50 mM). Data represents five technical replicates (n = 3). **F.** HIF-1α protein and mRNA levels were measured in LPS-stimulated GM-CSF macrophages under indicated sugar conditions (n = 3–4). **G–J.** SKG mice were injected with curdlan to induce arthritis and administered fructose in drinking water starting 3 days prior to immunization. Glucose levels in serum and synovial fluid (SF) were assessed at day 42 (n = 5–8). **H–I.** mRNA levels of Hif1a and HIF-1α target genes in synovial cells were analyzed by RT-qPCR (n = 5). **J.** Intracellular HIF-1α expression was measured by flow cytometry in CD45⁺CD11b⁺F4/80⁺ synovial cells (n = 9–10). Graphs represent mean ± SEM. \**p* < 0.05; \*\**p* < 0.01; \*\*\**p* < 0.001; \*\*\*\**p* < 0.0001 by Mann–Whitney U test (A–C, E–J) or unpaired t-test (D, F).

IL-1β production requires two sequential steps: the synthesis of intracellular pro-IL-1β and its proteolytic cleavage by caspase-1 into mature IL-1β^34^. In fructose-treated macrophages, IL-1β mRNA expression and pro-IL-1β protein levels were both increased, whereas caspase-1 activity remained unchanged (Fig. 2B-C). These results indicate that fructose enhances IL-1β production primarily through transcriptional regulation rather than post-translational processing in GM-CSF macrophages. Glycolysis-mediated stabilization of HIF-1α is a key mechanism controlling IL-1β expression in macrophages^35^. Consistent with this, fructose treatment promoted glycolytic activity to a degree comparable to glucose, although the maximal glycolytic capacity was modestly reduced (Fig. 2D-E). Furthermore, HIF-1α protein levels were elevated following fructose exposure (Fig. 2F). These findings suggest that fructose supports glycolytic metabolism, stabilizes HIF-1α, and thereby potentiates downstream inflammatory signaling pathways in GM-CSF macrophages.

Chronic inflammatory environments, such as the inflamed synovial fluid in RA, are characterized by localized glucose depletion^36, 37^. Consistent with this, the synovial fluid of SKG mice exhibited lower glucose concentrations than serum, which was not altered by fructose supplementation. Thus, in this metabolic context fructose-derived glycolysis may compensate for limited glucose availability to sustain inflammatory activation (Fig. 2G). Indeed, SKG mice maintained on a fructose-enriched diet displayed a significant increase in Hif1a mRNA expression and upregulation of HIF-1α target genes in synovial cells (Fig. 2H–I), indicating enhanced activation of HIF-1α–dependent transcriptional programs. Among synovial cell populations, macrophages showed particularly robust elevation of HIF-1α expression in response to fructose (Fig. 2J). These results demonstrate that under glucose-limited conditions, fructose serves as an alternative metabolic substrate that promotes glycolysis, stabilizes HIF-1α, and amplifies inflammatory gene expression in GM-CSF macrophages, thereby contributing to sustained inflammatory responses within arthritic tissues.

### Fructose supports hexokinase (HK)-dependent glycolysis in inflammatory macrophages

To elucidate the metabolic role of fructose in inflammatory macrophages, a targeted metabolomics analysis was performed on LPS-stimulated GM-CSF macrophages cultured with either fructose or glucose. This analysis focused on key intermediates of the glycolytic pathway. While both sugars enhanced glycolytic metabolite production relative to carbohydrate-free controls (Fig. 3A), glucose-treated macrophages exhibited a more substantial accumulation of intermediates between glucose-6-phosphate (G6P) and glyceraldehyde-3-phosphate (G3P) compared with fructose-treated cells. Despite these intermediate-level differences, lactate concentrations, representing the endpoint of glycolysis, were comparable between the two groups (Fig. 3B), indicating similar overall glycolytic flux. Notably, the NAD⁺/NADH ratio was significantly lower in glucose-treated macrophages than in those treated with fructose (Supplementary Fig. 3A). Because G3P is metabolized by glyceraldehyde-3-phosphate dehydrogenase (GAPDH), which utilizes NAD⁺ as a cofactor (Supplementary Fig. 3B)^38^, the reduced NAD⁺/NADH ratio likely suppressed GAPDH activity, leading to accumulation of upstream glycolytic intermediates in glucose-treated cells (Fig. 3A). Together, these data indicate that fructose can sustain glycolytic flux in GM-CSF macrophages similarly to glucose, but with a distinct redox profile.

**Figure 3.**
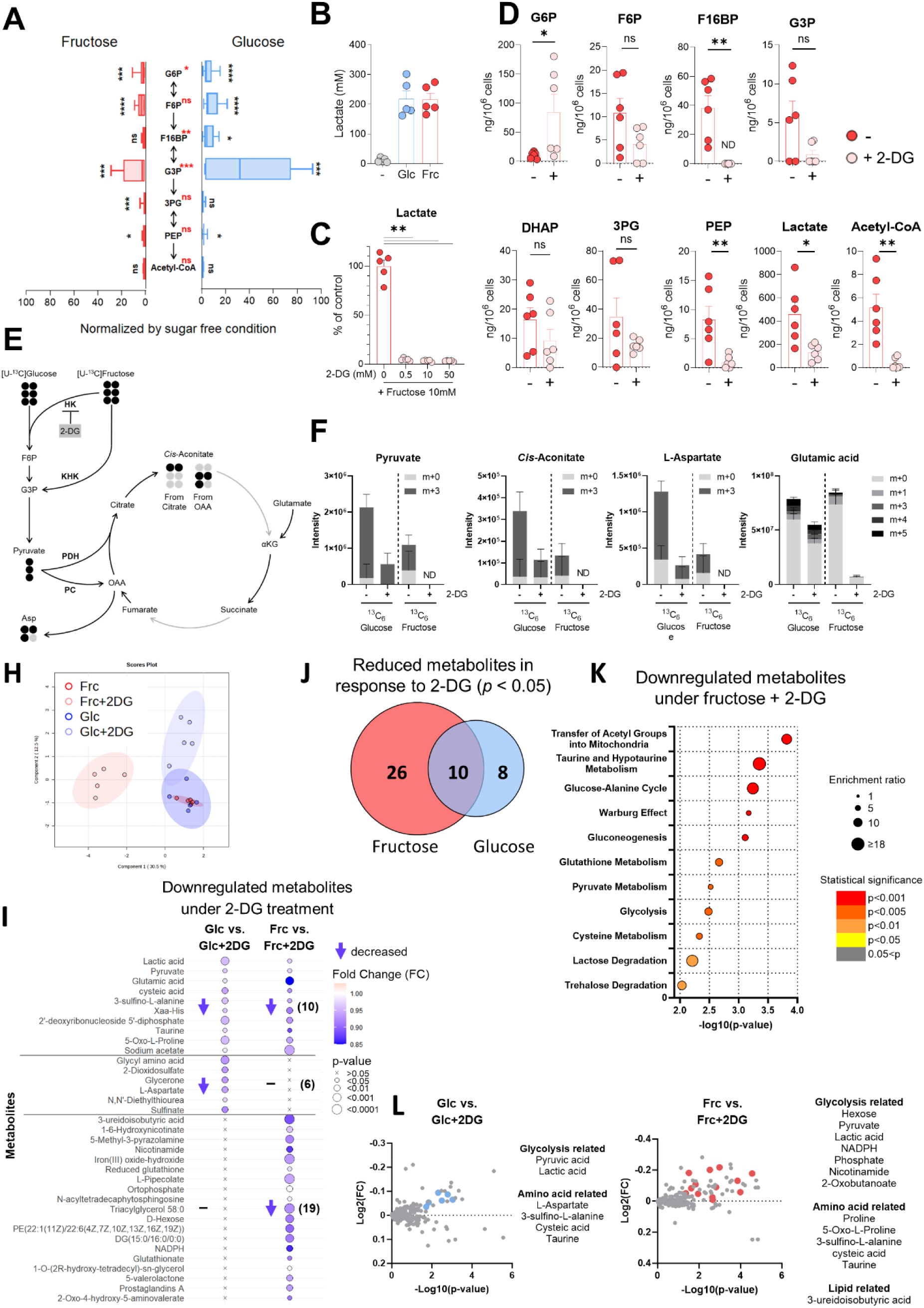
Fructose undergoes glycolysis *via* hexokinase (HK)-dependent metabolism in human macrophages. Human GM-CSF macrophages were pre-starved of glucose for 2 h and then stimulated with LPS (50 ng/mL) for 18 h in glucose and serum-free medium supplemented with 10 mM glucose or fructose. **A.** LC-MS analysis of intracellular glycolytic intermediates [fructose-6-phosphate (F6P), fructose-1,6-bisphosphate (F16BP), 3-phosphoglyceric acid (3PG), and phosphoenolpyruvic acid (PEP)] under indicated conditions. **B–C.** Lactate concentrations in culture supernatants were measured after supplementation with sugars or 2-DG. **D.** LC-MS quantification of glycolytic intermediates following 2-DG (0.5 mM) treatment in fructose-supplemented macrophages. Each dot indicates an individual sample analyzed**. E.** Schematic of stable isotope tracing experiment with ^13^C-labeled glucose or fructose. Dark and light circles represent ^13^C and ^12^C atoms, respectively (HK; Hexokinase, 2-DG; 2-deoxyglucose, KHK; Ketohexokinase, G3P; glyceraldehyde-3-phosphate, PDH; pyruvate dehydrogenase, PC; pyruvate carboxylase, OAA; oxaloacetic acid, αKG; alpha-ketoglutarate). **F.** Relative intensities of labeled and unlabeled metabolic products (pyruvate, cis-aconitate, aspartate, glutamate) were quantified. **G.** PLS-DA plot demonstrates distinct intracellular metabolomic profiles among four groups: glucose/fructose ± 2-DG. (Cross-validation: Accuracy = 0.9, R² = 0.9687, Q² = 0.5800). **H–K.** Metabolomic comparisons reveal downregulated metabolites with 2-DG treatment. Dot plot indicates fold-change and significance of metabolites. The numbers in parentheses indicate the number of metabolites (H). Venn diagram displays metabolite reduction in response to 2-DG (I). Pathway enrichment analysis of downregulated metabolites under fructose + 2-DG (J). Volcano plots illustrate metabolomic changes after 2-DG under glucose or fructose conditions; Blue and red dots represent the metabolites that were significantly hit in pathway enrichment analysis (K). Graphs show mean ± SEM. \**p* < 0.05; \*\**p* < 0.01; \*\*\**p* < 0.001; \*\*\*\**p* < 0.0001 by Mann–Whitney U test.

Fructose can enter glycolysis through two distinct enzymatic routes: phosphorylation by hexokinase (HK) to form fructose-6-phosphate (F6P), or phosphorylation by ketohexokinase (KHK) to generate glyceraldehyde-3-phosphate (G3P)^39^. Because the conversion of F6P to fructose-1,6-bisphosphate (F1,6BP) is an irreversible step, fructose-derived G6P and F6P must proceed through the HK-dependent glycolytic pathway for continued metabolism. To assess this dependency, we employed 2-deoxyglucose (2-DG), a competitive inhibitor of HK. Treatment with 0.5 mM 2-DG completely abolished fructose-induced lactate production (Fig. 3C), whereas glycolysis in glucose-treated macrophages was only partially inhibited at the same concentration, consistent with glucose’s higher affinity for HK (Supplementary Fig. 3C)^40^. These results demonstrate that fructose-driven glycolysis in inflammatory macrophages is critically dependent on HK activity (Fig. 3D).

To further confirm fructose incorporation into central carbon metabolism, ¹³C metabolic flux analysis was performed using ¹³C₆-labeled fructose or glucose (Fig. 3E). ¹³C from fructose was incorporated into glycolytic intermediates, including pyruvate, as well as into tricarboxylic acid (TCA) cycle metabolites such as *cis*-aconitate and aspartate (Fig. 3F). In agreement with prior observations (Fig. 3A and 3D), fructose contributed to central carbon metabolism, albeit less efficiently than glucose. Importantly, 0.5 mM 2-DG completely blocked ¹³C incorporation from fructose, reaffirming that HK is indispensable for fructose metabolism in GM-CSF macrophages.

To comprehensively assess the metabolic impact of glucose and fructose beyond glycolysis, untargeted metabolomics profiling was conducted in GM-CSF macrophages (Supplementary Fig. 4A). To delineate HK-dependent metabolic effects, 2-DG was applied under both sugar conditions. This analysis revealed that 2-DG treatment caused substantial shifts in metabolite profiles, particularly in fructose-treated macrophages (Fig. 3G). Correlation analyses demonstrated considerable overlap between glucose- and fructose-treated conditions; however, a distinct pattern of metabolic reprogramming emerged specifically under fructose plus 2-DG treatment (Fig. 3G; Supplementary Fig. 4B). Differential metabolite analysis under 2-DG inhibition identified significant alterations in various metabolites across diverse metabolic pathways, including glycolysis and amino acid metabolism (Fig. 3H; Supplementary Fig. 4C). Among the significantly decreased metabolites, 10 metabolites shared similar profile changes in both sugar conditions, whereas 6 and 19 metabolites displayed sugar-dependent alterations unique to glucose and fructose, respectively (Fig. 3H). Conversely, among the 31 significantly increased metabolites, 12 and 10 were sugar-dependent, and 9 metabolites showed opposite trends between glucose and fructose conditions (Supplementary Fig. 4C). Notably, a greater number of metabolites were responsive to 2-DG in fructose-treated macrophages (50 in total) than in glucose-treated macrophages (35 in total) (Fig. 3I; Supplementary Fig. 4D), indicating that HK inhibition exerts a more pronounced impact on intracellular metabolism when fructose is the primary carbon source.

Metabolite Set Enrichment Analysis (MSEA) of differentially expressed metabolites (Fig. 3J; Supplementary Fig. 4E–G) corroborated that 2-DG significantly perturbs glycolysis and associated metabolic networks, with more pronounced effects in fructose-treated macrophages. This was further supported by comparison of the number of significantly altered metabolic pathways in each condition, where both the absolute set size and intersection size were markedly greater among decreased metabolites under fructose treatment. Metabolites decreased by 2-DG in fructose-treated macrophages relative to other conditions (Supplementary Fig. 4H) were largely associated with glycolysis and amino acid metabolism (Fig. 3K). Collectively, these findings underscore the heightened sensitivity of fructose-driven metabolic pathways to HK inhibition and demonstrate that fructose metabolism in inflammatory GM-CSF macrophages is tightly governed by HK-dependent glycolytic flux.

### KHK-dependent fructose metabolism promotes the formation of methylglyoxal-derived advanced glycation end-products (MG-AGEs) in inflammatory macrophages

Given the distinct metabolomic profiles observed upon fructose treatment (Fig. 3G–K), we hypothesized that fructose activates metabolic pathways distinct from those engaged by glucose. Ketohexokinase (KHK) and aldolase B (ALDOB), the key enzymes of the fructolytic pathway, are primarily expressed in the liver and brain^41^. Notably, during macrophage differentiation, both KHK and ALDOB expression levels increased, with ALDOB showing particularly strong induction in GM-CSF-differentiated macrophages (Fig. 4A).

**Figure 4.**
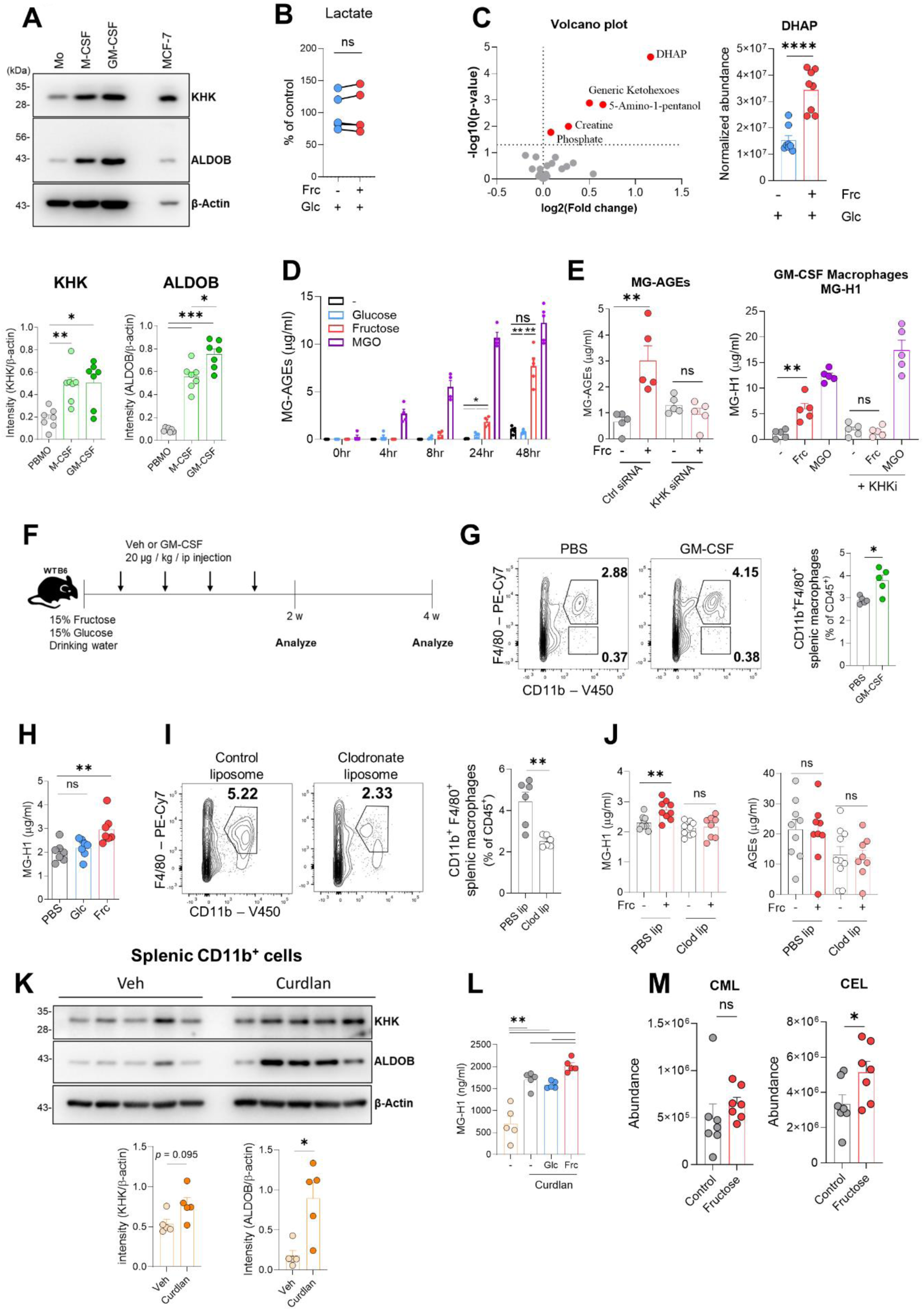
Fructolysis promotes MG-AGE formation in inflammatory macrophages. **A.** Protein expression of KHK and ALDOB was assessed in human monocytes cultured with M-CSF or GM-CSF for 6 days. MCF-7 cells served as positive controls. **B-E.** Human GM-CSF macrophages were stabilized for 2 h and then cultured in complete medium supplemented with 50 ng/mL LPS for 24 h or indicated time in the presence or absence of additive 10 mM sugar. Lactate levels were quantified in LPS-stimulated GM-CSF macrophages cultured with or without sugars (B). Volcano plot shows differentially expressed metabolites post-fructose exposure; Normalized abundance of DHAP is highlighted (C). MG-H1 levels in supernatants were quantified over time after 10 mM glucose, fructose, or 0.5 mM MGO treatment (D). MG-H1 levels after siRNA-mediated knockdown of KHK in GM-CSF macrophages (E). **F.** Schematic of *in vivo* experimental design in WT mice receiving GM-CSF and fructose or glucose in drinking water for 4 weeks. **G.** Frequency of splenic CD45⁺CD11b⁺F4/80⁺ macrophages at week 2 post GM-CSF or vehicle (n = 5 per group). **H.** Serum MG-H1 levels at week 4 (n = 7 per group). **I–J.** After clodronate depletion, macrophage frequency and serum MG-H1 and AGE levels were evaluated (n = 6–10). **K.** KHK and ALDOB protein expression in CD11b⁺ splenic cells from SKG mice post-curdlan (n = 5). **L–M.** Serum MG-H1 and AGE levels measured by ELISA or LC-MS in SKG mice with fructose supplementation post-curdlan (n = 5–7). Graphs show mean ± SEM. \**p* < 0.05; \*\**p* < 0.01; \*\*\**p* < 0.001; \*\*\*\**p* < 0.0001 by Mann–Whitney *U* test.

To better approximate physiological conditions, fructose was supplemented into glucose-containing medium (10 mM glucose). Under these conditions, fructose did not alter lactate production from aerobic glycolysis but significantly increased intracellular levels of dihydroxyacetone phosphate (DHAP) (Supplementary Fig. 5A–C). ¹³C-tracing analysis revealed that inhibition of HK-dependent fructose metabolism using 0.5 mM 2-deoxyglucose (2-DG) reduced ¹³C incorporation into pyruvate but not into DHAP (Supplementary Fig. 5B), indicating that the accumulation of DHAP arises through HK-independent metabolic routes in GM-CSF macrophages.

DHAP derived from fructose metabolism can be directed toward glycolysis, lipogenesis, or the synthesis of methylglyoxal (MGO), a reactive precursor of advanced glycation end-products (AGEs) (Supplementary Fig. 5C)^42^. Although fructose had minimal effects on glycolytic or lipogenic intermediates (Fig. 4B; Supplementary Fig. 5D), the concentration of soluble MG-AGEs, particularly MG-H1, was markedly elevated in the supernatants of fructose-treated GM-CSF macrophages (Fig. 4C). *In vitro* assays using BSA or FBS demonstrated that MG-AGEs were generated exclusively in the presence of fructose, but not glucose, when either 100 µg/mL BSA or 10% FBS was included in the culture medium (Supplementary Fig. 5E; Fig. 4D)^43^. Moreover, siRNA-mediated silencing of KHK or pharmacological inhibition of its enzymatic activity significantly attenuated MG-AGE formation (Supplementary Fig. 5F; Fig. 4E), confirming that KHK-dependent fructolysis is essential for MG-AGE production in inflammatory macrophages.

To determine whether inflammatory macrophages contribute to MG-AGE formation *in vivo*, wild-type (WT) mice were administered GM-CSF intraperitoneally to expand the macrophage population (Fig. 4F)^44^. Two weeks post-treatment, there was a significant increase in splenic CD45⁺CD11b⁺F4/80⁺ macrophages (Fig. 4G). Following four weeks of fructose consumption, serum MG-AGE levels were significantly higher in these mice compared with control or glucose-fed groups (Fig. 4H). In contrast, other AGE species associated with hepatic glycolysis remained unchanged between glucose- and fructose-fed mice (Supplementary Fig. 5G). Furthermore, macrophage depletion using clodronate liposomes at two weeks post-GM-CSF injection markedly reduced serum MG-AGE concentrations (Fig. 4I–J)^45, 46^, supporting a key role for macrophage-mediated fructolysis in systemic MG-AGE production.

In the pathological context of autoimmune arthritis, elevated ALDOB expression was detected in splenic CD11b⁺ cells from SKG mice, a model in which GM-CSF is critical for disease development (Fig. 4K)^47^. Consistent with the *in vitro* findings, serum from fructose-fed SKG mice exhibited significantly higher MG-AGE levels compared with controls, while other AGE types remained unaltered (Fig. 4L). Although curdlan treatment increased both MG-AGEs and other AGE species (Supplementary Fig. 5H), the fructose-specific effect was most pronounced for Nε-carboxyethyl-lysine (CEL), a methylglyoxal-derived AGE product generated *via* fructolysis. In contrast, levels of Nε-carboxymethyl-lysine (CML), typically produced from glyoxal through oxidative pathways, were unaffected (Fig. 4M)^48^. These findings demonstrate that KHK-dependent fructolysis in inflammatory macrophages promotes the generation of methylglyoxal-derived AGEs, particularly CEL, under chronic inflammatory conditions. This mechanism provides a direct metabolic link between fructose utilization and the exacerbation of inflammatory pathology.

### Fructose-induced AGEs in macrophages amplify pro-inflammatory responses in immune cells via RAGE signaling

To elucidate the impact of fructose on macrophage function, we performed bulk RNA sequencing of GM-CSF macrophages following 24-hour fructose exposure. Transcription factor enrichment analysis (TRRUST) identified several upregulated transcriptional regulators, among which MMP9 ranked prominently based on cumulative transcription factor (TF)-gene interaction scores (Fig. 5A and B). Consistent with this, MMP9 secretion was significantly increased at 24-48 hours after fructose treatment, but not at earlier time points (Fig. 5C). MMP9, a matrix metalloproteinase that mediates extracellular matrix (ECM) degradation and facilitates immune cell infiltration, is known to be transcriptionally activated by RAGE signaling^43^. Given that fructose simultaneously elevated methylglyoxal-derived AGEs (MG-AGEs) in culture supernatants during the same timeframe (Fig. 4D), we hypothesized that MG-AGEs engage RAGE to induce MMP9 expression. Supporting this model, fructose treatment upregulated the expression of full-length RAGE (fl-RAGE), the functional membrane-bound receptor variant (Fig. 5D). MMP9 secretion was robustly induced by fructose, but not by glucose, and was significantly attenuated upon ketohexokinase (KHK) inhibition, implicating KHK-dependent fructolysis in MMP9 upregulation (Fig. 5E and F). Moreover, pharmacological blockade of RAGE signaling suppressed MMP9 expression (Fig. 5G), confirming that the MG-AGE–RAGE axis mediates this effect.

**Figure 5.**
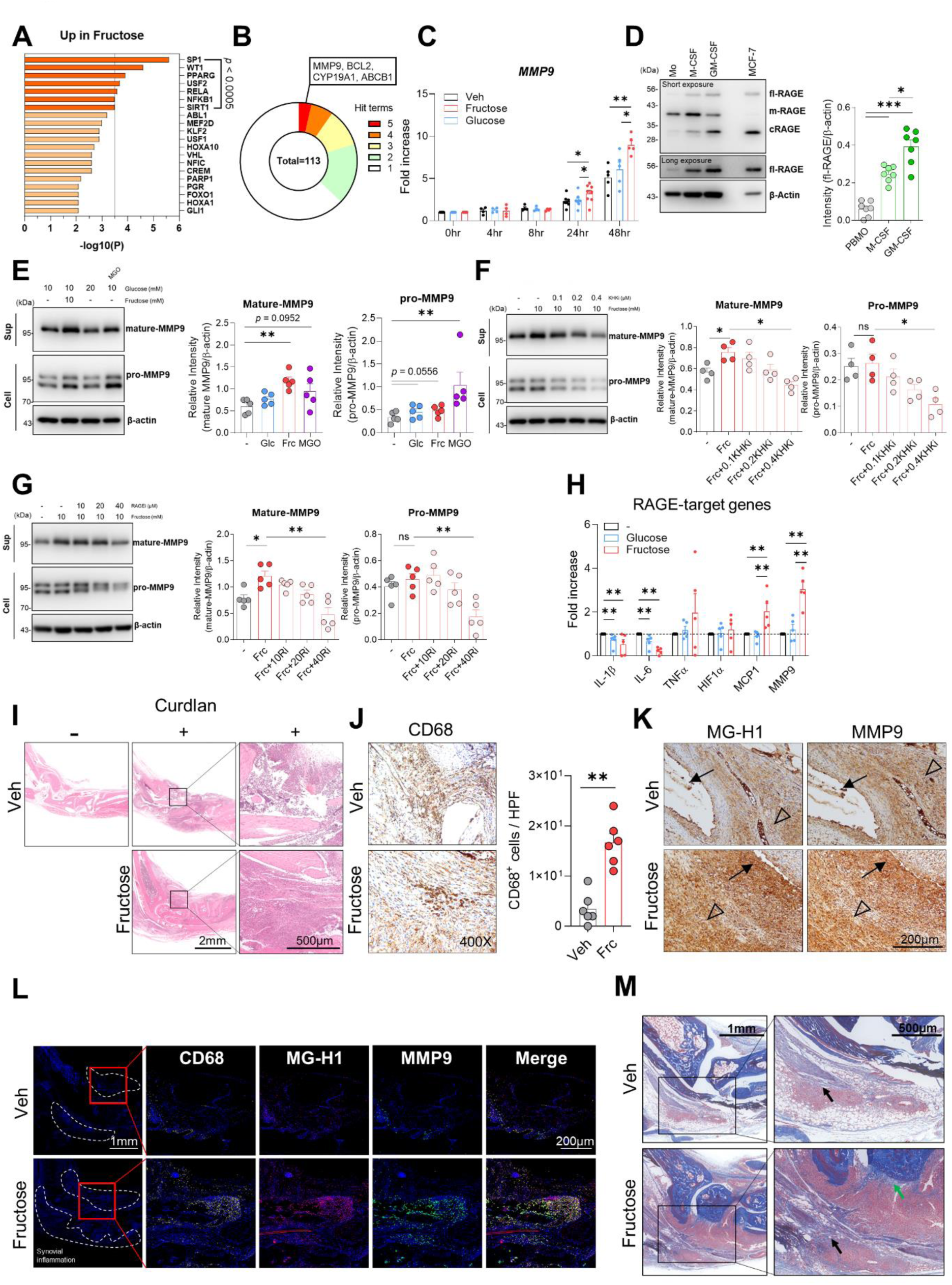
Fructose-derived MG-AGEs promote macrophage inflammation *via* RAGE signaling. **A–B.** DEGs from LPS-stimulated GM-CSF macrophages with/without fructose (10 mM) were analyzed using TRRUST *via* Metascape (*p*-value < 0.3, LogFC > 0.15) (A). Key transcription factor networks are shown (n = 3) (B). **C.** MMP9 mRNA expression of human GM-CSF macrophages was measured in response to glucose or fructose (10 mM) post-LPS stimulation (n = 4–8). **D.** RAGE protein levels were analyzed in M-CSF and GM-CSF differentiated macrophages (n = 7); MCF-7 cells served as control. **E–G.** Human GM-CSF macrophages were stimulated for 36 h with LPS in the presence of sugar, 2-DG, or MGO used as a positive control to mimic MG-AGE-driven responses. **H.** mRNA levels of IL1B, IL6, TNFA, HIF1A, MMP9, and MCP1 were measured after treatment with conditioned media from glucose- or fructose-treated macrophages (n = 5). **I–M.** Fructose was administered to SKG mice in drinking water from 3 days before curdlan injection until sacrifice. H&E-stained paw sections reveal synovial inflammation (n = 3) (I). CD68 immunostaining shows increased macrophage infiltration in fructose-treated mice (n = 6) (J). **K–L.** Immunostaining for MG-H1 and MMP9 in synovial tissue, with multiplex immunofluorescent staining highlighting colocalization of CD68, MG-H1, and MMP9 (n = 3–6). **M.** M-T staining reveals cartilage (green arrow) and collagen fibrosis (black arrow) in the tarsus area of fructose-treated SKG mice (n = 3). Graphs show mean ± SEM. \**p* < 0.05; \*\**p* < 0.01; \*\*\**p* < 0.001 by Mann–Whitney U test.

To determine whether secreted MG-AGEs directly propagate inflammatory signaling, we transferred supernatants from fructose-treated macrophages to freshly differentiated GM-CSF macrophages. Within 8 hours, recipient cells exhibited elevated expression of canonical RAGE target genes, including MMP9 and MCP-1, while expression of TNF, HIF-1α, IL-1β, and IL-6 remained unchanged (Fig. 5H). These findings indicate that fructose-derived MG-AGEs act as soluble mediators that activate RAGE in an autocrine and paracrine manner to promote selective pro-inflammatory gene expression. *In vivo*, fructose-fed SKG mice exhibited enhanced immune cell infiltration within arthritic joints (Fig. 5I) and increased accumulation of CD68⁺ macrophages (Fig. 5J). Concordantly, fructose administration induced strong upregulation of MMP9 and MG-AGEs within ankle joint tissues (Fig. 5K). Multiplex immunofluorescence confirmed co-localization of MG-H1 and MMP9 within CD68⁺ synovial myeloid cells in fructose-fed mice (Fig. 5L). Furthermore, Masson’s trichrome staining revealed pronounced ECM degradation in fructose-treated joints (Fig. 5M). These findings demonstrate that fructose-induced MG-AGEs secreted by inflammatory macrophages activate RAGE signaling, leading to enhanced MMP9 expression and ECM remodeling. This autocrine/paracrine MG-AGE–RAGE loop amplifies inflammatory responses and may contribute to tissue destruction in chronic inflammatory disorders such as autoimmune arthritis.

### Macrophage-derived MG-AGEs regulate CD4⁺ T cell differentiation via RAGE Signaling

Chronic inflammatory diseases, including autoimmune disorders, are characterized by complex crosstalk between innate and adaptive immune compartments^49^. Given that fructose-fed mice exhibited heightened proinflammatory activity in both macrophages and CD4⁺ T cells (Fig. 1), we next sought to determine whether fructose directly modulates T-cell responses. We first analyzed the expression of fructose metabolism-related genes following TCR stimulation. GLUT5 (SLC2A5) expression was upregulated in naive CD4⁺ T cells upon TCR activation, whereas KHK expression remained unchanged (Fig. 6A). In a T cell-only culture system, fructose supplementation did not alter the differentiation of Th1 or Th17 cells, regardless of glucose availability (Fig. 6B), suggesting that fructose alone is insufficient to modulate T-cell polarization. In contrast, a co-culture system of naive CD4⁺ T cells with monocytes, stimulated with anti-CD3/CD28 and lipopolysaccharide (LPS), respectively (Fig. 6C), revealed distinct metabolic and immunological outcomes. LPS stimulation progressively upregulated GLUT5 (SLC2A5) expression in monocytes, and fructose supplementation led to a substantial increase in methylglyoxal-derived AGEs (MG-AGEs) in the culture supernatant by day 6 (Fig. 6D and E). Under these conditions, production of IL-17A and IFN-γ was elevated (Fig. 6F), indicating that fructose enhances monocyte-dependent promotion of proinflammatory T cell differentiation.

**Figure 6.**
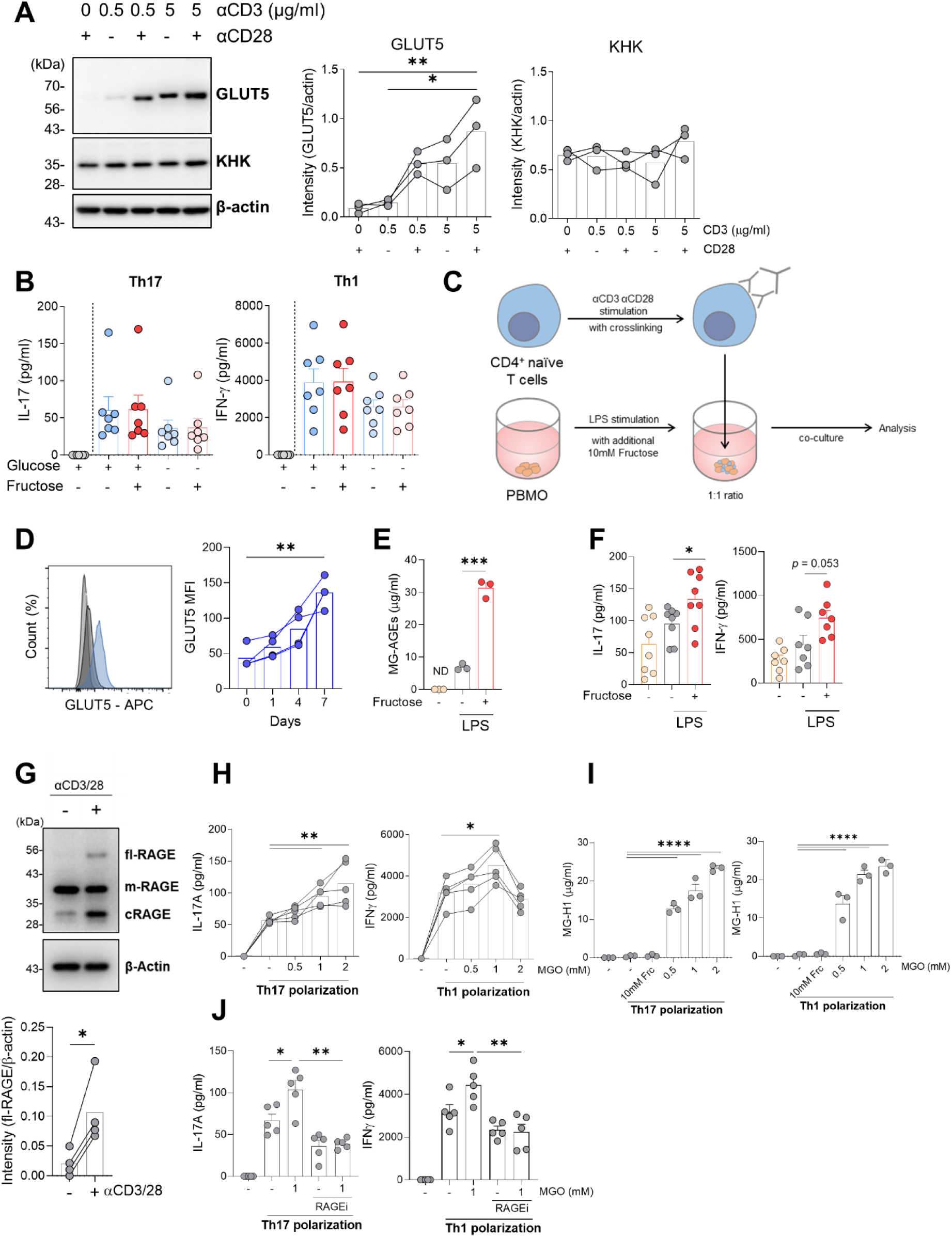
Fructose modulates GM-CSF macrophage function and influences naive CD4⁺ T-cell differentiation. **A.** Protein expression of GLUT5 and KHK was assessed in PBMC-derived naive CD4⁺ T cells following CD3/CD28 stimulation by immunoblotting (n = 4). Representative blot shown. **B.** Levels of IL-17A and IFN-γ were measured by ELISA from culture supernatants of Th17- or Th1-polarized naive CD4⁺ T cells treated with fructose or glucose for 7 days (n = 7). **C–F.** Naive CD4⁺ T cells were co-cultured with monocytes to evaluate the impact of monocyte-derived factors on T cell function. (C) Schematic representation of the co-culture system. (D) Surface expression of GLUT5 in CD14⁺ monocytes during LPS stimulation was assessed by flow cytometry (n = 4). (E) MG-AGE (MG-H1) levels were quantified in culture supernatants on day 6 of co-culture (n = 3). (F) IL-17A and IFN-γ levels were quantified in culture supernatants at day 13 (n = 7–8). **G.** RAGE protein expression in CD3/CD28-stimulated naive CD4⁺ T cells was analyzed by immunoblotting (n = 4). **H–I.** Naive CD4⁺ T cells were treated with methylglyoxal (MGO) and cultured under Th17 or Th1 conditions. (H) IL-17A and IFN-γ secretion was measured after 7 days by ELISA (n = 5). (I) MG-H1 levels in supernatants were assessed under the same conditions (n = 3). **J.** To assess the role of RAGE, naive CD4⁺ T cells were co-treated with MGO and a RAGE inhibitor (RAGEi) during Th17/Th1 polarization; IL-17A and IFN-γ were measured by ELISA (n = 5). Densitometry was used to quantify immunoblots normalized to β-actin. Data are presented as mean ± SEM. \**p* < 0.05, \*\**p* < 0.01, \*\*\**p* < 0.001, \*\*\*\**p* < 0.0001 by Mann–Whitney *U* test, one-way ANOVA (D, I), or unpaired *t*-test (E).

Furthermore, TCR stimulation induced full-length (fl) RAGE expression in naive CD4⁺ T cells (Fig. 6G). Exposure to methylglyoxal (MGO, 1 mM) during Th1 and Th17 polarization augmented their differentiation, correlating with dose-dependent increases in MG-H1, a representative MG-AGE, in the supernatant (Fig. 6H and I). Importantly, pharmacological inhibition of RAGE suppressed MGO-induced Th1 and Th17 differentiation (Fig. 6J), confirming that MG-AGE–RAGE signaling mediates this effect. These findings suggest that MG-AGEs generated by macrophages promote CD4⁺ T cell differentiation through RAGE-dependent signaling. This supports the existence of a fructose-macrophage-T cell axis, in which fructose-derived MG-AGEs amplify inflammatory responses by both inducing MMP9-mediated matrix remodeling and enhancing proinflammatory T cell polarization *via* MG-AGE–RAGE signaling, thereby exacerbating tissue inflammation.

### Fructose metabolism in macrophages is associated with human RA pathology

To evaluate the clinical relevance of our experimental findings, we examined whether the fructolytic characteristics observed in GM-CSF macrophages are reflected in monocytes/macrophages from patients with rheumatoid arthritis (RA). Given that GM-CSF levels are elevated in the peripheral blood of RA patients, we compared the fructose responsiveness of CD14⁺ peripheral blood monocytes (PBMOs) from healthy controls (HC) and RA patients. When cultured in glucose-depleted medium and stimulated with LPS, RA PBMOs produced significantly higher levels of lactate in response to fructose than did HC PBMOs, indicating enhanced fructose-driven glycolytic activity in RA (Fig. 7A). Similarly, in complete medium supplemented with fructose, RA PBMOs exhibited robust accumulation of the fructose-derived AGE MG-H1, whereas HC PBMOs produced only negligible amounts under any condition. This effect was abolished by KHK inhibition, while glucose supplementation alone had no impact, supporting that KHK-dependent fructolysis is functionally active in RA PBMOs even under glucose-replete conditions (Fig. 7B).

**Figure 7.**
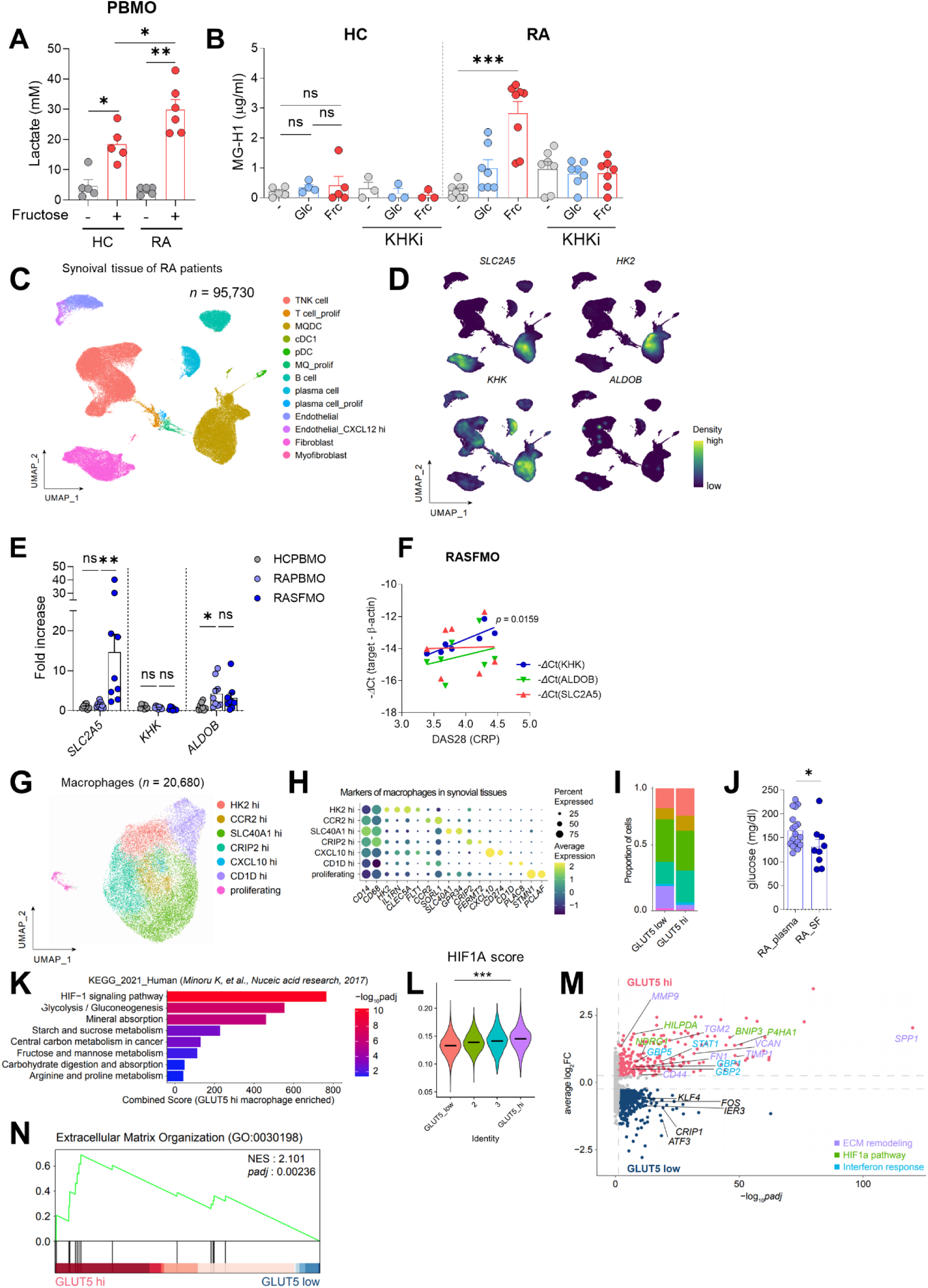
Fructose metabolism is associated with macrophage inflammatory responses in RA patients. **A.** Human monocytes were glucose-starved for 2 h and cultured for 18 h in glucose-free, serum-free medium containing 50 ng/mL LPS with or without 10 mM fructose. Lactate concentrations in culture supernatants were quantified. **B.** Monocytes were pre-stabilized for 2 h, then cultured in complete medium with 50 ng/mL LPS for 48 h in the presence or absence of 10 mM sugar or a KHK inhibitor. Soluble MG-AGEs (MG-H1) were measured in supernatants. **C.** Uniform manifold approximation and projection (UMAP) analysis of human RA synovial tissue defined major immune cell types. **D.** Expression levels of SLC2A5, KHK, HK2, and ALDOB across identified clusters. **E.** RT-qPCR quantification of fructolysis and glycolysis-related genes (SLC2A5, KHK, and ALDOB) in monocytes from healthy PBMCs (HC PBMO), RA PBMCs (RA PBMO), and RA synovial fluid monocytes (RA SFMO). **F.** Correlation analysis between KHK, ALDOB, and SLC2A5 expression and disease activity (DAS28-CRP) in RA SFMO. **G.** UMAP visualization of macrophage subsets annotated by fine-grained transcriptional states. **H.** Dot plot showing the expression of marker genes across macrophage clusters. **I.** Bar plot indicating relative proportions of GLUT5^hi^ and GLUT5^low^ macrophages. **J.** Glucose levels measured in RA plasma and synovial fluid. **K.** Bar plots of enriched KEGG pathways from differentially expressed genes (DEGs) between GLUT5^hi^ and GLUT5^low^ macrophages, ranked by combined score (odds ratio × adjusted *P* value). **L.** Violin plots of HIF1α module scores across macrophage clusters stratified by SLC2A5 expression. **M.** Volcano plot of DEGs between GLUT5^hi^ and GLUT5^low^ macrophages; ECM remodeling (purple), HIF1α signaling (green), and interferon response (blue) genes are highlighted. **N.** Gene set enrichment analysis (GSEA) for “Extracellular Matrix Organization” (GO:0030198), enriched in GLUT5^hi^ macrophages. NES, normalized enrichment score. Data represent mean ± SEM. \**p* < 0.05, \*\**p* < 0.01, \*\*\**p* < 0.001, \*\*\*\**p* < 0.0001 (Mann–Whitney *U* test or Kruskal-wallis test).

To further investigate fructose metabolism at the site of inflammation, we reanalyzed publicly available single-cell RNA sequencing (scRNA-seq) data from RA synovial tissue (syn52297840)^50^. Synovial macrophages demonstrated pronounced expression of SLC2A5, KHK, and HK2, suggesting enhanced fructolytic capacity (Fig. 7C–D, Supplementary Fig. 6A–B). These findings were validated by RT-qPCR, which revealed significant upregulation of SLC2A5 and ALDOB in RA synovial fluid monocytes (SFMOs), and elevated ALDOB expression in RA PBMOs compared with controls (Fig. 7E). Although group-level differences in KHK expression were not observed, KHK transcript levels positively correlated with DAS28(CRP) scores in RA SFMO, indicating that fructolytic potential scales with inflammatory disease activity (Fig. 7F).

Consistent with these data, scRNA-seq analysis confirmed that CD14⁺CD68⁺ synovial cells, primarily macrophages, expressed SLC2A5, KHK, and HK2 across subsets (Fig. 7G–H, Supplementary Fig. 6C–D). Using a Gaussian mixture model, we classified macrophages into four populations (1; GLUT5^lo^, 2-3; GLUT5^mid^, 4; GLUT5^hi^) (Supplementary Fig. 6E). Notably, GLUT5^hi^ macrophages exhibited elevated expression of inflammatory mediators such as S100A9, SERPINA1, and CD44 compared with GLUT5^lo^ counterparts (Supplementary Fig. 6F).

Functionally, GLUT5^hi^ macrophages contained a markedly higher proportion of HK2 ^hi^ cells than did GLUT5^lo^ macrophages (Fig. 7I). As expected, glucose concentrations were significantly lower in RA synovial fluid than in matched plasma samples (Fig. 7J), indicating that monocytes within the joint encounter a nutrient-restricted, hypoglycemic environment. Pathway enrichment analysis demonstrated that GLUT5^hi^ macrophages were enriched for HIF-1 signaling and glycolysis-related pathways, accompanied by elevated HIF1A activity scores (Fig. 7K–L). Moreover, this subset displayed increased expression of extracellular matrix (ECM) remodeling–related genes, including MMP9, suggesting a coupling between fructose metabolism and tissue-destructive programs (Fig. 7M–N). These findings identify fructose metabolism as a distinct pathogenic feature of RA myeloid cells. Through KHK-dependent fructolysis, GLUT5^hi^ macrophages promote MG-AGE formation, inflammatory activation, and ECM remodeling, thereby contributing to joint inflammation and structural damage in RA.

## DISCUSSION

For most of human history, human exposure to fructose was minimal and intermittent, limited to seasonal intake from fruits and honey. The advent of industrial food production and the widespread use of high-fructose corn syrup (HFCS) have dramatically altered this nutritional landscape. In contrast to glucose, whose metabolism is tightly regulated by insulin signaling and key rate-limiting steps in glycolysis (such as phosphofructokinase-1), fructose is metabolized via ketohexokinase (KHK) and enters the glycolytic pathway downstream of these regulatory checkpoints^51^. This comparatively unregulated entry into cellular metabolism promotes lipogenesis and oxidative stress, supporting the view that sustained high fructose intake constitutes an evolutionary mismatch between contemporary dietary patterns and the mechanisms of human metabolic regulation^52^.

Epidemiological and experimental data increasingly implicate chronic fructose consumption in the development of inflammatory diseases. Long-term fructose feeding in mice induces non-alcoholic fatty liver disease (NAFLD) and hepatic inflammation, primarily through activation of Kupffer cells and accumulation of lipid intermediates^53, 54^. Notably, emerging evidence indicates that fructose can exert immunomodulatory effects independent of overt metabolic syndrome^24, 25, 55, 56, 57, 58^. For example, epidemiological analyses have revealed an association between high fructose intake and increased arthritis prevalence even among non-obese individuals^59^. Consistent with these observations, our study shows that a six-week 15% fructose diet markedly exacerbated arthritis in SKG mice in the absence of hepatic steatosis, suggesting that fructose can potentiate inflammation independently of hepatic dysfunction (Fig. 1H–N; Supplementary Figure 1). These findings support the notion that fructose modulates immune responses locally, particularly within macrophage populations, thereby implicating fructose as a broader regulator of immune function beyond its established role in metabolic disease. In the present study, we identify a previously unrecognized dual fructose metabolic program in proinflammatory macrophages that sustains inflammation via both HK-and KHK-dependent pathways and contributes to the pathogenesis of autoimmune disease *in vitro* and *in vivo*.

A key contribution of this work is the identification of macrophage fructose metabolism as a central driver of inflammatory adaptation under conditions of nutrient stress (Fig. 3 and 4). Inflamed tissues such as the rheumatoid synovium represent nutrient-deprived microenvironments characterized by hypoxia, low glucose availability, and lactate accumulation^60, 61, 62, 63, 64^. Because macrophages rely heavily on glycolysis to support their bioenergetic and effector functions^65^, glucose scarcity poses a major constraint on the maintenance of inflammation. Here, we demonstrate that GM-CSF-differentiated macrophages, which resemble proinflammatory tissue-resident macrophages, acquire the capacity to use fructose as an alternative carbon source, thereby preserving glycolytic flux when glucose is limited. This metabolic flexibility is driven by GM-CSF-induced expression of the fructose transporter GLUT5 and the rate-limiting enzyme KHK (Fig. 1F and 4A; Supplementary Figure 1D). GLUT5 expression is transcriptionally regulated by c-Myc, as pharmacologic inhibition of c-Myc abrogates GLUT5 induction (Supplementary Figure 1C), confirming its essential role in enabling macrophage fructose utilization. These observations are consistent with previous reports that c-Myc coordinates the expression of glucose and amino acid transporters and reinforce its function as a metabolic rheostat that allows immune cells to adapt to nutrient stress^30, 31, 32^. By facilitating fructose utilization, GM-CSF effectively rewires macrophage metabolism to support survival and activation under glucose-limited conditions, extending prior findings that GM-CSF enhances glycolysis in myeloid cells by revealing fructose metabolism as a parallel compensatory pathway (Fig. 3). This mechanism is likely to be particularly relevant within chronically inflamed microenvironments such as the rheumatoid joint (Fig. 1I–N).

Jones and colleagues recently reported that fructose reprograms cellular metabolism to favor glutaminolysis and oxidative phosphorylation, thereby supporting enhanced inflammatory cytokine production in LPS-stimulated human monocytes^24^. In line with these findings, our data indicate that circulating monocytes express only low levels of GLUT5, KHK, and ALDOB, which are required for canonical KHK-dependent fructolysis (Fig. 1F and Fig. 4A. Notably, we observed that GLUT2, a transporter capable of mediating uptake of both glucose and fructose, is highly expressed in peripheral blood monocytes but is markedly downregulated during macrophage differentiation (Supplementary Figure 7). This differential expression pattern provides a plausible explanation for the distinct fructose-driven metabolic programs observed between human monocytes and GM-CSF-differentiated macrophages, and suggests that lineage- and differentiation-dependent transporter regulation critically shapes how myeloid cells exploit fructose under inflammatory conditions.

Our study further demonstrates that macrophage fructose metabolism generates specific intermediates that propagate inflammation. KHK-mediated fructolysis promotes the accumulation of dihydroxyacetone phosphate (DHAP), which serves as a precursor for methylglyoxal (MGO) production (Fig. 4C)^66^. MGO is a highly reactive α-dicarbonyl that non-enzymatically modifies amino acid residues to form methylglyoxal-derived advanced glycation end-products (MG-AGEs), including Nε-(carboxyethyl)lysine (CEL) and MG-H1. In contrast to glucose-derived AGEs such as carboxymethyl lysine (CML), which arise predominantly through oxidative pathways, MG-AGE formation is tightly associated with heightened glycolytic and fructolytic flux^67, 68, 69, 70^. We show that short-term fructose exposure selectively increases MG-H1 accumulation in macrophages, independently of hepatic dysfunction or systemic metabolic syndrome (Fig. 4E). These findings identify macrophages as a previously underappreciated source of MG-AGEs under fructose-replete conditions and implicate KHK-dependent fructolysis as a local driver of carbonyl stress. Crucially, MG-H1 accumulation was related with heightened arthritis severity in SKG mice, establishing a mechanistic link between macrophage fructose metabolism, MG-AGE–mediated carbonyl stress, and tissue inflammation (Fig. 4L).

The pathogenic consequences of MG-AGEs are mediated *via* the receptor for advanced glycation end-products (RAGE), which is abundantly expressed on macrophages and other immune cells^71, 72, 73^. Engagement of RAGE by MG-AGEs triggers oxidative stress and downstream signaling cascades that perpetuate inflammation^71, 73, 74, 75^. Notably, fructose-treated macrophages exhibited marked upregulation of MMP9 in a RAGE-dependent manner, whereas canonical cytokines such as IL-1β and IL-6 were minimally affected (Fig. 5A-H). This selective activation suggests that MG-AGE–RAGE signaling preferentially drives extracellular matrix remodeling and leukocyte migration, processes central to joint destruction in rheumatoid arthritis^43, 76, 77^. Given MMP9’s established role in matrix degradation and tissue infiltration, its induction by fructose-derived AGEs provides a direct mechanistic link between altered metabolism and structural tissue damage.

Transcriptomic and metabolic analyses of rheumatoid arthritis synovial macrophages support this model (Fig. 7). Synovial macrophages exhibited increased expression of SLC2A5, KHK, and HK2, together with upregulation of MMP9, HIF1A, and glycolytic enzymes. Consistent with this metabolic phenotype, glucose concentrations in synovial fluid were substantially lower than in plasma, reinforcing the concept of nutrient limitation that favors fructose utilization. Moreover, monocytes from RA patients produced increased lactate and MG-H1 upon fructose exposure, whereas these effects were abolished by KHK inhibition (Fig. 7B), demonstrating active KHK-dependent fructose metabolism in human disease. Collectively, these findings identify fructose utilization as a defining feature of proinflammatory macrophages within nutrient-limited inflammatory niches.

Together, our data establish a mechanistic framework in which GM-CSF–induced metabolic reprogramming enables macrophages to exploit fructose to sustain energy demands under glycolytic restriction. While this adaptation supports cellular survival, it concurrently fuels the accumulation of MGO and MG-AGEs, activating RAGE signaling and promoting MMP9-mediated tissue remodeling and Th17 polarization. This self-reinforcing loop links dietary fructose intake to macrophage-driven inflammation and tissue damage, providing a metabolic explanation for the exacerbation of joint pathology in fructose-fed mice.

Although characterized in arthritis, this mechanism likely extends to other macrophage-driven pathologies. Tissue-resident macrophages (TRMs) in the liver, adipose tissue, brain, and tumor microenvironments exhibit metabolic flexibility and are therefore likely susceptible to fructose-driven inflammation^78^. Kupffer cells exposed to high portal fructose flux contribute to NAFLD progression^53^, adipose macrophages activate RAGE signaling under metabolic stress to promote insulin resistance^79^, and microglial RAGE activation disrupts the blood–brain barrier and accelerates neuroinflammation in Alzheimer’s disease^74, 80, 81^. Similarly, tumor-associated macrophages utilize MMP9-dependent matrix remodeling to facilitate angiogenesis and metastasis^82, 83^. We propose that fructose-driven MG-AGE accumulation may serve as a shared metabolic driver across these conditions, linking modern dietary patterns to chronic inflammation and tissue remodeling.

In summary, our findings redefine fructose as an active immunometabolic regulator rather than a passive nutrient. Through GM-CSF–c-Myc–dependent reprogramming, macrophages acquire the capacity to metabolize fructose under glucose-limited conditions, ensuring survival at the cost of increased carbonyl stress and RAGE activation. This metabolic adaptation initiates a self-perpetuating inflammatory cascade that promotes tissue damage and sustains chronic inflammation. Therapeutically, targeting key nodes of this pathway such as KHK, RAGE, or c-Myc, may offer new strategies to disrupt fructose-driven immune activation. As global fructose consumption continues to rise, understanding how fructose reprograms immune cell metabolism will be essential for developing interventions that mitigate the impact of modern dietary excess on chronic inflammatory disease.

## METHODS

### Preparation of human PBMCs and SFMCs

Study protocols were reviewed and approved by the IRB (Institutional Review Board) of Seoul National University Hospital and Chungnam National University Hospital. Peripheral blood was drawn after obtaining written, informed consent. Characteristics of RA patients enrolled in this study are summarized in Supplementary Tables 1 and 2. Collected synovial fluid or peripheral blood were processed within 2 h of fluid or plasma collection. Glucose levels were immediately measured using a Glucolab glucometer (Osang Healthcare, Anyang, Republic of Korea) following the manufacturer’s instructions. Peripheral blood mononuclear cells (PBMCs) or synovial fluid mononuclear cells (SFMCs) were isolated by density gradient centrifugation (Biocoll separating solution; BIOCHROM Inc., Cambridge, UK).

### Isolation and culture of human monocytes

Monocytes were positively separated from PBMC or SFMC with anti-CD14 microbeads (Miltenyi Biotec Inc., Auburn, CA, USA). Purified monocytes were cultured in RPMI 1640 medium (Welgene, Kyungsan, Republic of Korea) supplemented with 10% fetal bovine serum or glucose depleted RPMI 1640 medium (Thermo Fisher Scientific, Waltham, MA, USA) with 1% penicillin/streptomycin and 1% l-glutamine. To obtain monocyte-derived macrophages, purified CD14^+^ monocytes were cultured in the presence of recombinant human GM-CSF or M-CSF (50 ng/ml of each; PeproTech, Rocky Hill, NJ, USA) for 6 days. Cells were stimulated with LPS (Thermo Fisher Scientific) in the presence of the indicated chemical inhibitors or reagents, including ATP (Sigma-Aldrich, St. Louis, MO, USA), 2-DG (Sigma-Aldrich), BSA (Bovogen Biologicals, Melbourne, VIC), methylglyoxal (MGO; MilliporeSigma, Burlington, MA, USA), KHKi (MedKoo Biosciences Inc., Morrisville, NC, USA), and RAGEi (MilliporeSigma). All sugars (D-(+)-glucose, D-(-)-fructose) for replenishing experiments were purchased from Sigma-Aldrich.

### Isolation and culture of human CD4⁺ T Cells

Naive CD4⁺ T cells were isolated from human peripheral blood mononuclear cells (PBMCs) via negative selection using the MojoSort™ Human CD4⁺ Naive T Cell Isolation Kit (BioLegend, San Diego, CA) following the manufacturer’s instructions. Purified cells were cultured in RPMI 1640 medium supplemented with 10% fetal bovine serum (FBS), 1% penicillin/streptomycin, and 1% L-glutamine (hereafter, complete RPMI 1640). For T cell activation, cells were stimulated with anti-CD3/CD28-coated Dynabeads (T-Activator CD3/CD28; Thermo Fisher Scientific, Waltham, MA) at a bead-to-cell ratio of 1:10 in the presence or absence of the indicated reagents, including methylglyoxal (MGO; MilliporeSigma). To induce Th17 or Th1 polarization, naïve CD4⁺ T cells were cultured in either serum-free X-VIVO 10 media (Lonza, Basel, Switzerland) for Th17 differentiation or in complete RPMI 1640 for Th1 differentiation. Cells were plated in 96-well U-bottom plates and stimulated with anti-CD3/CD28-coated beads for 7 days under the following cytokine conditions: recombinant human (rh) IL-1β (30 ng/mL), rhIL-6 (30 ng/mL), rhIL-23 (10 ng/mL), and rhTGF-β (10 ng/mL) (all from R&D Systems, Minneapolis, MN) for Th17 polarization, rhIL-2 (100 IU/mL; PeproTech) and rhIL-12 (10 ng/mL; R&D Systems) for Th1 polarization, and rhIL-2 (100 IU/mL) alone for non-polarizing control. For co-culture experiments, TCR-stimulated naive CD4⁺ T cells were cultured with LPS-activated monocytes at a 1:1 ratio, as previously described^84, 85^. Culture supernatants were collected on day 6 or day 13, and cytokine concentrations were measured by ELISA.

### Quantitative RT-PCR

Total RNA was extracted using TRIzol reagent (Life Technologies) and reverse-transcribed using the GoScript™ system (Promega). Quantitative real-time PCR was performed using the Bio-Rad CFX system with SYBR Green (Bio-Rad). Primers were designed using Primer-BLAST or obtained from published sequences (listed in Supplementary Table 3). Expression levels were normalized to ACTINB using the *ΔΔ*Ct method.

### Immunoblot analysis

Total proteins were prepared using radioimmunoprecipitation assay (RIPA) buffer (150 mM NaCl, 10 mM Na2HPO4, pH 7.2, 1% Nonidet P-40, and 0.5% deoxycholate) containing PMSF (phenylmethylsulfonyl fluoride; Millipore Sigma), EDTA, and a protease and phosphatase inhibitor cocktail (Thermo Fisher Scientific). Cell lysates were separated on an 8-10% SDS-polyacrylamide gel and blotted onto a polyvinylidene difluoride (PVDF) membrane (Bio-Rad) followed by blocking for 1 h with 5% BSA in Tris-buffered saline solution (TBS) containing 0.1% Tween 20. Membranes were incubated overnight at 4 °C with primary Abs, such as anti-GLUT2, anti-RAGE, anti-MMP9, and anti-KHK (all from Abcam, Cambridge, UK), anti-ALDOB (Thermo Fisher Scientific), anti-GLUT5 (Invitrogen, Waltham, MA, USA), and anti-HIF-1α (Cell Signaling Technology, Danvers, MA) Abs, followed by incubation with the HRP-conjugated secondary Ab for 1 h at RT. Membranes were developed using SuperSignal West Femto Maximum Sensitivity substrate or SuperSignal West Pico PLUS Chemiluminescent Substrate system (Thermo Fisher Scientific).

### Enzyme-linked immunosorbent assay (ELISA) and AGE quantification

The amounts of IL-17A, IFN-γ, IL-1β, IL-6, IL-23, GM-CSF, M-CSF, MG-H1, and AGEs in culture supernatant, plasma, or serum were quantified using commercially available human ELISA kits according to the manufacturer’s instructions (human IL-17A, IL-1β, IL-6, IL-23, GM-CSF, M-CSF, and mouse IL-17 ELISA kits from Invitrogen, Human IFN-γ ELISA MAX Deluxe from BioLegend, San Diego, CA, and MG-H1 and AGEs ELISA kits from Cell Biolabs Inc., San Diego, CA, USA). Measurement of OD (optical density) was performed using Infinite M200 Pro Multimode microplate reader (Tecan, Männedorf, Switzerland).

### Cell Viability

Lactate dehydrogenase (LDH) release was measured using the CytoTox 96® assay kit (Promega) to assess cell viability, following the manufacturer’s protocol.

### Flow cytometric analysis

Cultured cells or cells isolated from tissue were stained for 10 min with 7AAD (BD Biosciences, San Jose, CA, USA) to exclude dead cells. To analyze surface transporters and cells populations, cells were stained at 4°C for 30 min with Abs to GLUT5 (R&D Systems), CD45, CD11b (two from BD Bioscience), and F4/80 (Invitrogen). To analyze the expression of HIF-1α, isolated mouse immune cells were stained at 4°C for 30 min with Abs to CD45, CD11b (all from BD Bioscience), and F4/80 (Invitrogen), followed by fixation and permeabilization using Fix/Perm buffer (BioLegend). The fixed cells were stained with Abs against HIF-1α (R&D Systems). Stained cells were assessed using a BD LSRFortessa (BD Bioscience) and analyzed using FlowJo software (Tree Star, Ashland, OR).

### Mouse models of autoimmune arthritis and macrophage activation

Female SKG mice (8 weeks) used for induction of autoimmune arthritis were purchased from CLEA Japan (Tokyo, Japan). All mice were housed and maintained in a pathogen-free facility at Seoul National University (SNU) College of Medicine. All experiments were approved by the SNU Institutional Animal Care and Use Committee (Permission ID: SNU-230825-2-2). Mice were injected intraperitoneally with 3 mg/mice curdlan (FUJIFILM Wako Pure Chemical Corporation, Osaka, Japan) in 300 μl sterile PBS at 8 weeks. Ankle thicknesses were measured using calipers (Manostat, New York, NY). Joint swelling was monitored in blinded cages by two independent observers and scored as described (14647385): 0, no joint swelling; 0.1, swelling of one finger joint; 0.5, mild swelling of wrist, ankle, or base of tail; and 1.0, severe swelling of wrist, ankle or base of tail. Scores for all joints were totaled for each mouse. Mice were euthanized at the end of the experiment and immune cells were prepared from the spleen, inguinal LNs, and joints as previously described^86^. Tissues were collected for flow cytometry, RNA extraction, and histology. For GM-CSF-driven *in vivo* macrophage activation, wild-type C57BL/6 mice were injected intraperitoneally with recombinant mouse GM-CSF (PeproTech, 20 μg/kg, two times per week) and sacrificed after 2 or 4 weeks. Macrophage depletion was performed by intraperitoneal injection of clodronate liposomes (Liposoma B.V., Amsterdam, The Netherlands) at 100 μL/mouse twice a week for 2 weeks following GM-CSF injection^87^.

### Histology and immunohistochemistry

Ankle joints were fixed in 10% formalin, decalcified, embedded in paraffin, and sectioned. Formalin-fixed paraffin-embedded tissue blocks were cut into 4 μm-thick slices. Staining was performed with Masson’s Trichrome (Sigma) and immunohistochemistry using antibodies against MG-H1 (1:10000; Cell Biolabs), CD68 (1:16000), and MMP9 (1:5000; Abcam) and a Benchmark XT autostainer (Ventana Medical Systems, Tuscon, AZ), in accordance with the manufacturer’s instruction. Visualization was achieved using DAB substrate (Vector Labs). To examine histological alterations in the liver, the left lobe was fixed in 10% formalin and embedded in paraffin. Sections were prepared from the liver tissue blocks and stained with PAS, hematoxylin and eosin, and Masson’s Trichrome. Two expert pathologists reviewed all the specimens independently. Histological analyses of liver inflammation were performed by scoring based on steatosis, lobular inflammation ballooning, and fibrosis based on Supplementary Table 4.

### Multiplex Immunofluorescence

Multiplex immunofluorescence (IF) staining was performed on formalin-fixed, paraffin-embedded (FFPE) ankle sections using the Leica BOND RXm automated platform with the Opal 6-plex detection kit (Akoya, NEL871001KT) according to the provided 3-plex workflow. Sections (4 μm) were slide-baked (60 min) followed by heat-induced epitope retrieval (ER2, pH 9; 20 min). After washing with BOND Wash Solution 1, endogenous peroxidase was quenched and non-specific binding blocked with PKI Blocking Buffer (5 min). Primary antibodies were applied in three sequential cycles (Open1/2/3, 15 min each), each followed by Opal Polymer HRP (10 min) and the corresponding Opal fluorophore incubation (10 min): CD68 with Opal570 (1:1000), MMP9 with Opal520 (1:5000), and MG-H1 with Opal690 (1:20). Between cycles, antigen retrieval was performed with BOND ER Solution 1 at 95 °C for 20 min to remove bound antibodies while preserving tyramide deposits. All steps included intermediate washes with BOND Wash Solution 1. After the final cycle, nuclei were counterstained with Spectral DAPI (10 min), slides were washed and mounted. Stained sections were scanned and digitized using the PhenoImager HT 2.0 system (Akoya Biosciences) at 40× magnification.

### Metabolite extraction and preparation

Cells were washed twice with pre-warmed PBS (37 °C), detached on ice in PBS using a cell lifter, counted, and aliquoted at 1 × 10⁶ cells per pre-chilled tube. After centrifugation (2000 rpm, 5 min, 4 °C), pellets were resuspended in 500 μL ice-cold 50% acetonitrile, snap-frozen in liquid nitrogen, and stored at -80 °C. For extraction, samples were thawed on ice (15 min), vortexed (30 s), and sonicated (5 min, 4 °C), then centrifuged (15,000 rpm, 5 min, 4 °C). Supernatants were filtered through 0.2 μm syringe filters. For quantitative metabolomics, 10 μL of internal standard mix was added to 100 μL sample, vortexed (1 min, RT), centrifuged (14,000 rpm, 10 min, 4 °C), and 100 μL supernatant was transferred to vials; 5 μL was injected into the LC-MS/MS. For untargeted metabolomics, 10 μL from each sample was pooled for QC, while the remaining 90 μL was transferred to vials without internal standards; 4 μL was injected into the LC-Orbitrap-MS/MS.

### Quantitative analysis of glycolysis and TCA cycle metabolites

25 glycolytic and TCA intermediates (glucose, G6P, F6P, F16BP, DHAP, G3P, 3PG, PEP, acetyl-CoA, citrate, isocitrate, α-ketoglutarate, succinyl-CoA, succinate, malate, lactate, glutamine, glutamate, methionine, SAM, NAD, NADH, AMP, ADP, ATP) were quantified using an in-house LC-MS/MS method. Authentic standards and isotope-labeled internal standards (ATP-¹³C₁₀, AMP-¹³C₁₀,¹⁵N₅; Sigma-Aldrich) were dissolved at 1-5 mg/mL in 20 mM HCl/50% acetonitrile. Calibration and QC samples were prepared by serial dilution of mixed stocks in 50% acetonitrile (ranges, MRM transitions, and collision energies in Supplementary Table 5). Chromatography was performed on an Atlantis Premier BEH Z-HILIC column (2.1 × 100 mm, 1.7 μm; Waters) with an Agilent 1260 LC coupled to a 6490 triple quadrupole (Agilent). Mobile phases were 15 mM ammonium bicarbonate in water (A, pH 9) and in 90% acetonitrile (B, pH 9). Gradient: 90% B (0-0.1 min), 50% B (3.5-3.6 min), 90% B (6.5-10.0 min). Flow rate: 0.4 mL/min; column 37 °C. Detection used positive/negative ESI with MRM; capillary voltage -4000 V, nozzle -500 V; nitrogen as nebulizer (30 psi), drying gas (14 L/min, 250 °C), and sheath gas (11 L/min, 350 °C). Transitions were optimized with authentic standards. Data were processed with MassHunter Workstation vB.08.02 (Agilent Technologies).

### Untargeted and ¹³C-labeled untargeted metabolomics

Untargeted profiling was performed on an Ultimate 3000 UHPLC system (Thermo Fisher Scientific) coupled to a Q Exactive Plus Orbitrap mass spectrometer (Thermo Fisher Scientific) using an in-house untargeted method. Separation used an ACQUITY UPLC HSS T3 column (2.1 × 100 mm, 1.8 μm; Waters) at 40 °C. Mobile phases were 0.1% formic acid in water (A) and methanol (B) with gradient: 5% B (0-0.5 min, 0.35 mL/min), 5-95% B (0.5-3.5 min), 95% B (3.5-9.5 min), 95% B at 0.35-0.40 mL/min (9.5-15 min), 95% to 5% B (16.5-17.5 min), 5% B (17.5-20 min). Full MS scans (m/z 67-1000) were acquired at 120,000 resolution; spray voltage 4000 V (positive) and 3000 V (negative). Pooled QCs were analyzed in data-dependent MS/MS with NCE 10-50.

Raw files were converted to mzXML (MS) and MGF (MS2) using ProteoWizard (doi:10.1038/nbt.2377) and processed with tidyMass (doi:10.1038/s41467-022-32155-w) and X13CMS (doi:10.1038/s41596-019-0167-1; 10.1021/ac403384n). For untargeted data, peak picking used 5 ppm tolerance and 10-60 s peak widths; missing values were imputed by KNN, normalized with PQN, and log₂-transformed. Annotation was performed with metID (doi:10.1093/bioinformatics/btab583) and massDatabase (doi:10.1093/bioinformatics/btac546) against curated MS2 (e.g., MONA, MassBank) and MS1 (ChEBI, GNPS, KEGG) databases. Significant level-2 features were manually validated by spectral comparison.

For ¹³C-labeled datasets, centWave (5 ppm, 5-20 s) was used to identify labeled peaks, which were matched to untargeted data for annotation. Processed metabolite lists were analyzed with MetaboAnalyst (doi:10.1093/nar/gkae253) for PLS-DA, heatmaps, and pathway enrichment. Differential metabolite dot plots and UpSet plots were generated in R; volcano plots were created in GraphPad Prism v10.0.0 (GraphPad Software, Boston, MA, USA).

### Seahorse Metabolic Assays

To profile metabolic state of the cells, human primary macrophages in the presence of 100 ng/ml LPS (Sigma-Aldrich) for 24 h were seeded as a monolayer onto XFe24 cell culture plates (Seahorse Bioscience, MA, USA). The culture media were replaced with XF assay media supplemented with l-glutamine (300 µg/ml) and incubated for 1 h in a non-CO2 incubator. Glucose or fructose (10 mM), oligomycin (2 µM), and 2-DG (50 mM, all from Sigma-Aldrich) were sequentially added into the cells during real-time measurements of extracellular acidification rate (ECAR) and OCR (Oxygen consumption rate) using an XFe24 analyzer (Seahorse Bioscience). Glycolysis parameters were calculated using an XF glycolysis stress test report generator program which was provided by the manufacturer (Seahorse Bioscience). Glycolysis and glycolytic capacity were calculated by subtracting ECAR after glucose treatment from ECAR before oligomycin and subtracting ECAR after oligomycin from ECAR before 2-DG treatment, respectively.

### Lactate and triglyceride quantification assays

Lactate and triglyceride levels were measured using respective colorimetric assay kits (BioVision Technologies, Milpitas, CA, USA) following the manufacturer’s protocols. Absorbance for both assays was measured at 570 nm using an Infinite 200 PRO multimode microplate reader (Tecan, Männedorf, Switzerland).

### Single-cell RNA sequencing and data processing

Single-cell RNA-seq analysis of synovial tissue from patients with rheumatoid arthritis (RA) was performed using Seurat v4.0 (Satija Lab, New York, NY, USA) implemented in R v4.3.0.1. After generating the feature–barcode matrix, cells expressing fewer than 200 genes or more than 10% mitochondrial genes were excluded to remove low-quality cells. Normalization and identification of highly variable genes were performed using standard Seurat workflows. Principal component analysis (PCA) was carried out on the highly variable gene set, and the first 30 principal components (PCs) were used both to construct a shared nearest-neighbor (SNN) graph with FindNeighbors() and to embed the dataset into two dimensions using uniform manifold approximation and projection (UMAP). Clustering was performed using FindClusters() with the same set of 30 PCs.

### Differential gene expression analysis

Differentially expressed genes (DEGs) between GLUT5^hi^ and GLUT5^lo^ macrophages were identified using the MAST model implemented in Seurat’s FindMarkers() function. Genes expressed in at least 10% of cells within a given group were included in the DEG analysis. Resulting DEGs were ranked by average log2 fold change and adjusted P values were calculated using the Benjamini–Hochberg method to correct for multiple testing.

### Statistical Analysis

Two-tailed unpaired *t* tests, Mann-Whitney *U* tests, one-way ANOVAs with Kruskal-Wallis tests, or two-way ANOVAs were used to analyze the data using Prism 9 software (GraphPad Software Inc., La Jolla, CA), as indicated in the figure legends. p values less than 0.05 were considered to indicate statistical significance.

## ACKNOWLEDGMENTS

The authors thank Jiyeon Jang (Seoul National University College of Medicine) for assisting in the recruitment of human subjects and the Core Lab, Clinical Trials Center, Seoul National University Hospital for drawing blood.

## AUTHOR CONTRIBUTIONS

Conceptualization: YJK, SC, JYC, W-WL

Methodology: YJK, SC, JK, WS, JWP, EYL, DHC, SWK, JKP, JSL

Investigation: YJK, SC, JK, WS, YL, JWK, HYK, DJ

Visualization: YJK, SC, JK, WS, YL

Funding acquisition: YJK, W-WL

Supervision: JYC, W-WL

Writing – original draft: YJK, SC, WS, W-WL

Writing – review & editing: YJK, W-WL

