## Supplementary material for "Fructose utilization by GM-CSF-differentiated macrophages aggravates autoimmune inflammation *via* MG-derived AGE–RAGE signaling": Supplemntal table 1-2 and figure 1-7

### SUPPLEMENTARY MATERIALS

**Supplementary Table 1. The characteristics of cohort 1 RA patients (Figure 1).**

| RA peripheral blood | | |
|---|---|---|
| Patient Characteristics (Value) | Age (years), means ± SD | 57.84 ± 8.81 |
| | Sex (female/male), N (%) | 24/8 (75.00/25.00) |
| | Disease duration, months | 5.48 ± 5.12 |
| | Rheumatoid factor, no. of positive (%) | 28/32 (87.50) |
| | Rheumatoid factor titer (IU/ml), means ± SD | 122.90 ± 87.70 |
| | Anti-citrullinated protein antibody, no. of positive (%) | 25/32 (78.13) |
| | Anti-citrullinated protein antibody titer (IU/ml), means ± SD | 357.88 ± 161.59 |
| | ESR (mm/h), means ± SD | 26.81 ± 23.98 |
| | CRP (mg/dl), means ± SD | 2.00 ± 6.17 |
| | DAS28 ESR, means ± SD | 3.36 ± 1.62 |
| | DAS28 CRP, means ± SD | 2.42 ± 1.29 |

**Supplementary Table 2. The characteristics of cohort 2 RA patients (Figure 7).**

| Index | Remission | Low | Moderate | High |
|---|---|---|---|---|
| DAS28-erythrocyte sedimentation rate | <2.6 | ≥2.6 and ≤3.2 | >3.2 and ≤5.1 | >5.1 |
| CDAI | ≤2.8 | >2.8 and ≤10 | >10 and ≤22 | >22 |
| SDAI | ≤3.3 | >3.3 and ≤11 | >11 and ≤26 | >26 |

*DAS, disease activity score; CDAI, clinical disease activity index; SDAI, simplified disease activity index.*

| RA peripheral blood | | |
|---|---|---|
| Patient Characteristics (Value) | Age (years), means ± SD | 55.00 ± 12.0 |
| | Sex (female/male), N (%) | 16/2 (88.89/11.11) |
| | DAS28 ESR, means ± SD | 3.98 ± 1.95 |

| RA synovial fluid | | |
|---|---|---|
| Patient Characteristics (Value) | Age (years), means ± SD | 62.75 ± 11.1 |
| | Sex (female/male), N (%) | 5/3 (62.50/37.50) |
| | DAS28 ESR, means ± SD | 4.46 ± 0.43 |
| | DAS28 CRP, means ± SD | 3.92 ± 0.40 |

**Supplementary Table 3. Primers for qPCR.**

| Gene Name | Direction | Primer sequence (5'-3') |
|---|---|---|
| human HIF1A | Forward | CCATTAGAAAGCAGTTCCGC |
| human HIF1A | Reverse | TGGGTAGGAGATGGAGATGC |
| human BACTIN | Forward | GGACTTCGAGCAAGAGATGG |
| human BACTIN | Reverse | AGCACTGTGTTGGCGTACAG |
| human SLC2A5 | Forward | CAAGAAAGCCCTACAGACGCT |
| human SLC2A5 | Reverse | AGTAGATAGCGTTGACGCCC |
| human CXCL10 | Forward | CAGCAGAGGAACCTCCAGTC |
| human CXCL10 | Reverse | CAAAATTGGCTTGCAGGAAT |
| human IL1B | Forward | CCACAGACCTTCCAGGAGAATG |
| human IL1B | Reverse | GTGCAGTTCAGTGATCGTACAGG |
| human IL6 | Forward | AGACAGCCACTCACCTCTTCAG |
| human IL6 | Reverse | TTCTGCCAGTGCCTCTTTGCTG |
| human TNFA | Forward | TGCTTGTTCCTCAGCCTCTT |
| human TNFA | Reverse | CAGAGGGCTGATTAGAGAGAG |
| human MCP1 | Forward | AGAATCACCAGCAGCAAGTGTCC |
| human MCP1 | Reverse | TCCTGAACCCACTTCTGCTTGG |
| human MMP9 | Forward | GCCACTACTGTGCCTTTGAGTC |
| human MMP9 | Reverse | CCCTCAGAGAATCGCCAGTACT |
| human CASP1 | Forward | GCTGAGGTTGACATCACAGGCA |
| human CASP1 | Reverse | TGCTGTCAGAGGTCTTGTGCTC |
| human CXCL3 | Forward | TTCACCTCAAGAACATCCAAAGTG |
| human CXCL3 | Reverse | TTCTTCCCATTCTTGAGTGTGGC |
| human KHK | Forward | AATGCCTCCGTCATCTTCAGCC |
| human KHK | Reverse | ACCTGCTCTCACACGATGCCAT |
| human ALDOB | Forward | AGCCTCGCTATCCAGGAAAACG |
| human ALDOB | Reverse | TGGCAGTGTTCCAGGTCATGGT |
| human HK1 | Forward | CTGCTGGTGAAAATCCGTAGTGG |
| human HK1 | Reverse | GTCCAAGAAGTCAGAGATGCAGG |
| human HK2 | Forward | GAGTTTGACCTGGATGTGGTTGC |
| human HK2 | Reverse | CCTCCATGTAGCAGGCATTGCT |
| mouse ifng | Forward | CAGCAACAGCAAGGCGAAAAAGG |
| mouse ifng | Reverse | TTTCCGCTTCCTGAGGCTGGAT |
| mouse tgfb1 | Forward | TGATACGCCTGAGTGGCTGTCT |
| mouse tgfb1 | Reverse | CACAAGAGCAGTGAGCGCTGAA |
| mouse tgfb3 | Forward | AAGCAGCGCTACATAGGTGGCA |
| mouse tgfb3 | Reverse | GGCTGAAAGGTGTGACATGGAC |

| | | |
|---|---|---|
| mouse vegfa | Forward | CTGCTGTAACGATGAAGCCCTG |
| mouse vegfa | Reverse | GCTGTAGGAAGCTCATCTCTCC |
| mouse fas | Forward | CTGCGATTCTCCTGGCTGTGAA |
| mouse fas | Reverse | CAACAACCATAGGCGATTTCTGG |
| mouse srebp1 | Forward | CGACTACATCCGCTTCTTGCAG |
| mouse srebp1 | Reverse | CCTCCATAGACACATCTGTGCC |
| mouse hif1a | Forward | CCTGCACTGAATCAAGAGGTTGC |
| mouse hif1a | Reverse | CCATCAGAAGGACTTGCTGGCT |
| mouse bactin | Forward | AGCCATGTACGTAGCCATCC |
| mouse bactin | Reverse | CTCTCAGCTGTGGTGGTGAA |
| mouse il1b | Forward | TGGACCTTCCAGGATGAGGACA |
| mouse il1b | Reverse | GTTCATCTCGGAGCCTGTAGTG |
| mouse il6 | Forward | TACCACTTCACAAGTCGGAGGC |
| mouse il6 | Reverse | CTGCAAGTGCATCATCGTTGTTC |
| mouse il23a | Forward | CATGCTAGCCTGGAACGCACAT |
| mouse il23a | Reverse | ACTGGCTGTTGTCCTTGAGTCC |

**Supplementary Table 4. Histological analyses of liver inflammation.**

| Item | Definition | Score |
|---|---|---|
| **Steatosis (grade)** | Low to medium power evaluation of parenchymal involvement by steatosis | |
| | < 5% | 0 |
| | 5%-33% | 1 |
| | > 33%-66% | 2 |
| | > 66% | 3 |
| **Lobular Inflammation** | Overall assessment of all inflammatory foci | |
| | No foci | 0 |
| | < 2 foci per 200 × field | 1 |
| | 2-4 foci per 200 × field | 2 |
| | > 4 foci per 200 × field | 3 |
| **Ballooning** | | |
| | None | 0 |
| | Few balloon cells | 1 |
| | Many cells/prominent ballooning | 2 |
| **Fibrosis stage** | | |
| | None | 0 |
| | Perisinusoidal or periportal | 1 |
| | Mild, Zone 3, perisinusoidal | 1A |
| | Moderate, Zone 3, perisinusoidal | 1B |
| | Portal/periportal | 1C |
| | Perisinusoidal and portal/periportal | 2 |
| | Bridging fibrosis | 3 |
| | Cirrhosis | 4 |

**Supplementary Table 5. Calibration range, precursor and product ions, and collision energy for targeted metabolites analyzed by LC–MS/MS.**

| Metabolites | Abbreviation | Range (ng/mL) | Precursor ion | Product ion | Collision energy |
|---|---|---|---|---|---|
| Glucose | Glc | 50-10000 | 178.9 | 89.1 | -4 |
| Glucose-6-phosphate | G6P | 2-400 | 259.0 | 97.1 | -15 |
| Fructose-6-phosphate | F6P | 2-400 | 259.0 | 97.0 | -15 |
| Fructose-1,6-bisphosphate | F16BP | 20-4000 | 339.0 | 97.0 | -20 |
| Dihydroxyacetone phosphate | DHAP | 5-1000 | 168.8 | 96.9 | -10 |
| Glyceraldehyde 3-phosphate | G3P | 5-1000 | 168.8 | 78.9 | -20 |
| 3-Phosphoglycerate | 3PG | 10-2000 | 184.9 | 97.1 | -13 |
| Phosphoenolpyruvate | PEP | 2-400 | 166.9 | 79.0 | -10 |
| Acetyl coenzyme A | Acetyl-CoA | 0.5-100 | 810.0 | 303.1 | 35 |
| Citrate | | 50-10000 | 190.9 | 87.1 | -13 |
| Isocitrate | | 10-2000 | 191.0 | 73.1 | -22 |
| a-Ketoglutarate | | 10-2000 | 145.0 | 101.1 | -8 |
| Succinate | | 10-2000 | 117.1 | 73.1 | -7 |
| Malate | | 5-1000 | 133.0 | 115.1 | -8 |
| Lactate | | 50-10000 | 89.1 | 42.9 | -10 |
| Glutamine | | 10-2000 | 147.1 | 84.0 | 15 |
| Glutamate | | 20-4000 | 148.1 | 84.0 | 15 |
| Methionine | | 1-200 | 150.1 | 56.1 | 15 |
| S-Adenosyl methionine | SAM | 0.5-100 | 399.1 | 250.1 | 20 |
| Succinyl coenzyme A | Succinyl-CoA | 0.2-40 | 868.2 | 361.1 | 35 |
| Nicotinamide adenine dinucleotide | NAD | 10-2000 | 664.1 | 524.1 | 20 |
| Nicotinamide adenine dinucleotide reduced | NADH | 2-400 | 666.1 | 514.1 | 25 |
| Adenosine monophosphate | AMP | 2-400 | 348.0 | 136.0 | 18 |
| Adenosine triphosphate | ATP | 50-10000 | 508.3 | 136.2 | 25 |
| Adenosine diphosphate | ADP | 10-2000 | 428.2 | 136.1 | 25 |
| Adenosine-13C10,15N5 5′-triphosphate | ATP-13C10 | 5000 | 518.1 | 141.2 | 30 |
| Adenosine-13C10,15N5 5'-monophosphate | AMP-13C10,15N5 | 5000 | 361.1 | 78.9 | -20 |

#### Supplementary Figure 1

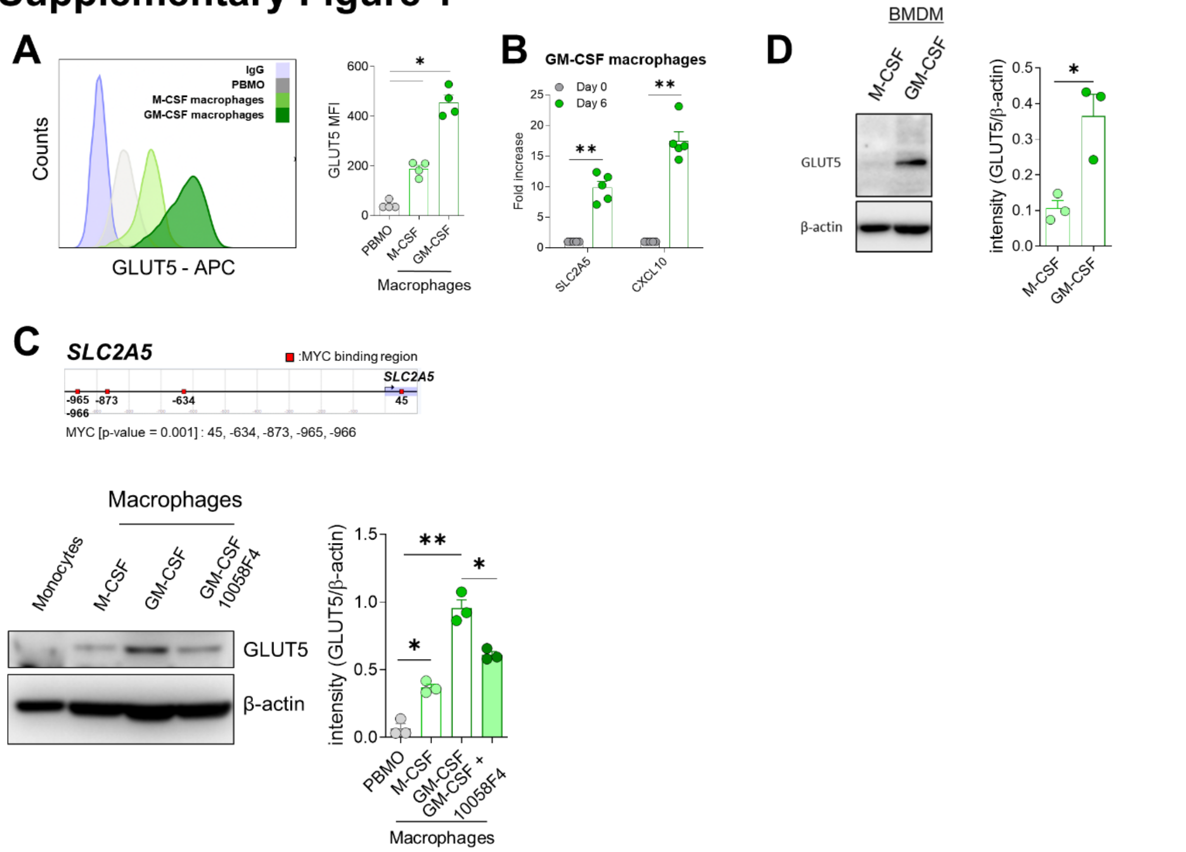


**Supplementary Figure 1. GM-CSF induces SLC2A5 (GLUT5) expression in human and murine macrophages. A.** Flow cytometric analysis of surface GLUT5 expression on CD14$^{+}$ monocytes from healthy controls (HCs) and on monocyte-derived macrophages differentiated with M-CSF (50 ng/mL) or GM-CSF (50 ng/mL) for 6 days. **B.** Time-course RT-qPCR analysis of SLC2A5 and CXCL10 mRNA expression in HC-derived monocytes cultured with GM-CSF (50 ng/mL) for the indicated durations. **C**. MYC motif analysis in the SLC2A5 promoter using the Eukaryotic Promoter Database (EPD) and the JASPAR CORE 2018 vertebrate motif library (upper). Representative immunoblot of GLUT5 protein expression in HC-derived monocytes differentiated with M-CSF (50 ng/mL) or GM-CSF (50 ng/mL) for 6 days (n = 3), pre-treated with the c-Myc inhibitor 10058-F4 for 30 min before GM-CSF stimulation (lower left). **D.** Murine bone marrow progenitor cells were isolated, differentiated into bone marrow–derived macrophages (BMDMs) using M-CSF or GM-CSF, and analyzed for GLUT5 protein expression on day 6. Representative immunoblots are shown (left). Band intensities were quantified by densitometry and normalized to β-actin. Graphs represent mean ± SEM. *$p$ < 0.05 and **$p$ < 0.01 by Mann–Whitney $U$ or unpaired $t$-test (A, B) or unpaired Student's t-test (C, D).

### Supplementary Figure 2

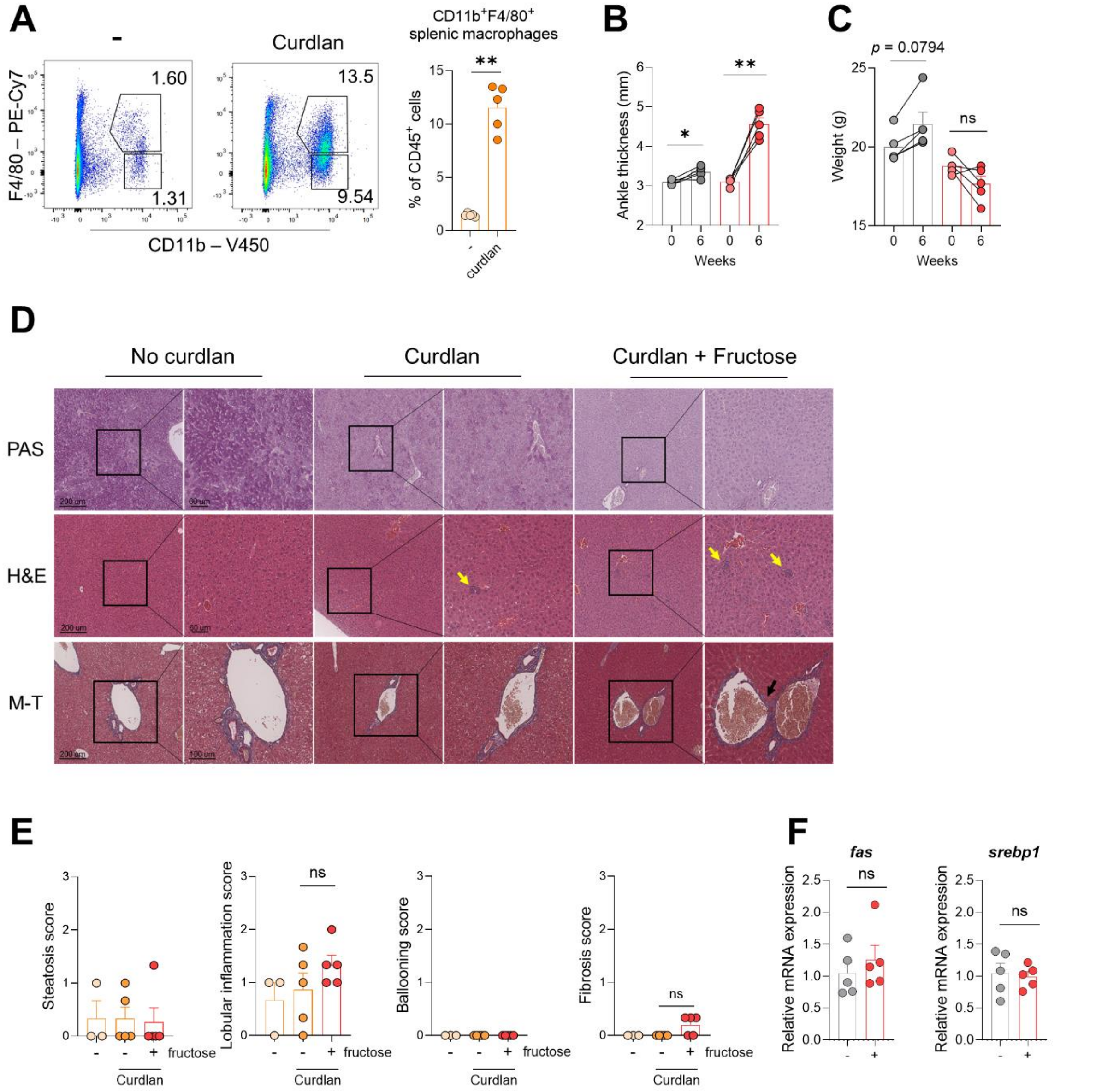


**Supplementary Figure 2. Fructose does not significantly affect hepatic inflammation in the SKG arthritis model. A.** Flow cytometric quantification of CD11b$^{+}$F4/80$^{+}$ macrophages among splenic CD45$^{+}$ cells from SKG mice six weeks after arthritis induction. **B-C.** Ankle thickness and body weight of SKG mice treated with or without fructose were monitored (n = 5 per group). **D.** Histological analysis of liver tissue (left lobe) at day 42 post-curdlan injection, stained with PAS, hematoxylin and eosin (H&E), and Masson's trichrome (M–T). Areas of lobular inflammation (yellow arrows, H&E) and fibrosis (black arrows, M–T) are indicated. **E.** Quantification of inflammatory area scores averaged from three representative sites (bottom, middle, top) of the left lobe. **F.** Relative hepatic mRNA expression of fas and srebp1 in arthritic mice (n = 5 per group). Graphs represent mean ± SEM. $*p < 0.05$ and $**p < 0.01$ by Mann–Whitney $U$ or unpaired $t$-test.

#### Supplementary Figure 3

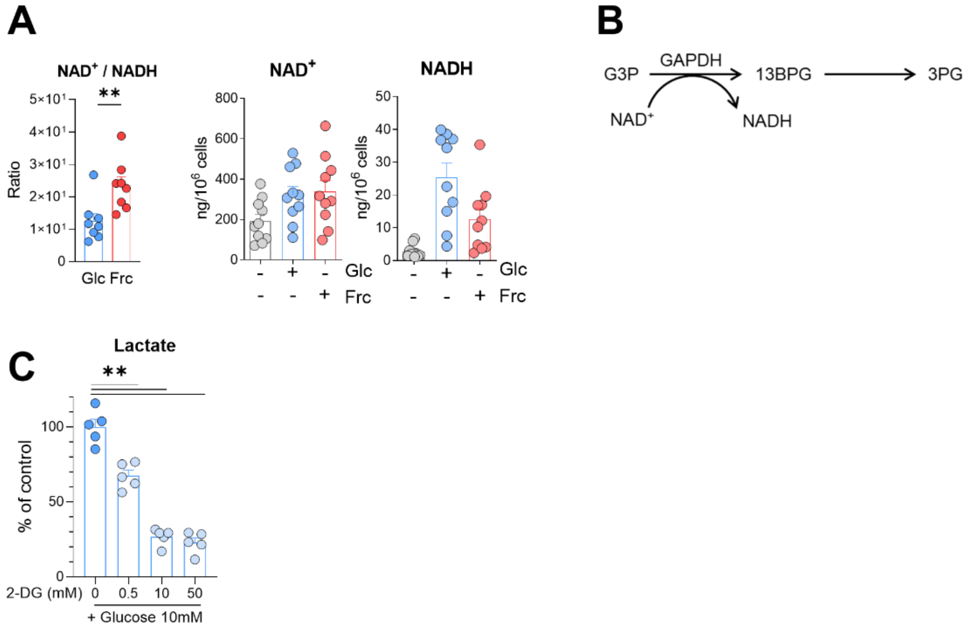


**Supplementary Figure 3. Fructose metabolism modulates the NAD$^+$/NADH balance in macrophages.** Human macrophages were pre-starved of glucose for 2 h, then cultured in glucose-depleted, serum-free medium containing 50 ng/mL LPS for 18 h with or without 10 mM glucose or fructose. **A.** LC–MS quantification of intracellular NAD$^+$ and NADH levels under the indicated conditions. **B.** Schematic overview of GAPDH-mediated glycolytic flux. **C.** Measurement of lactate concentrations in culture supernatants in the presence of increasing doses of 2-deoxyglucose (2-DG) combined with 10 mM glucose. Data represent mean ± SEM. $^{**}p < 0.01$ by Mann–Whitney *U* test.

### Supplementary Figure 4

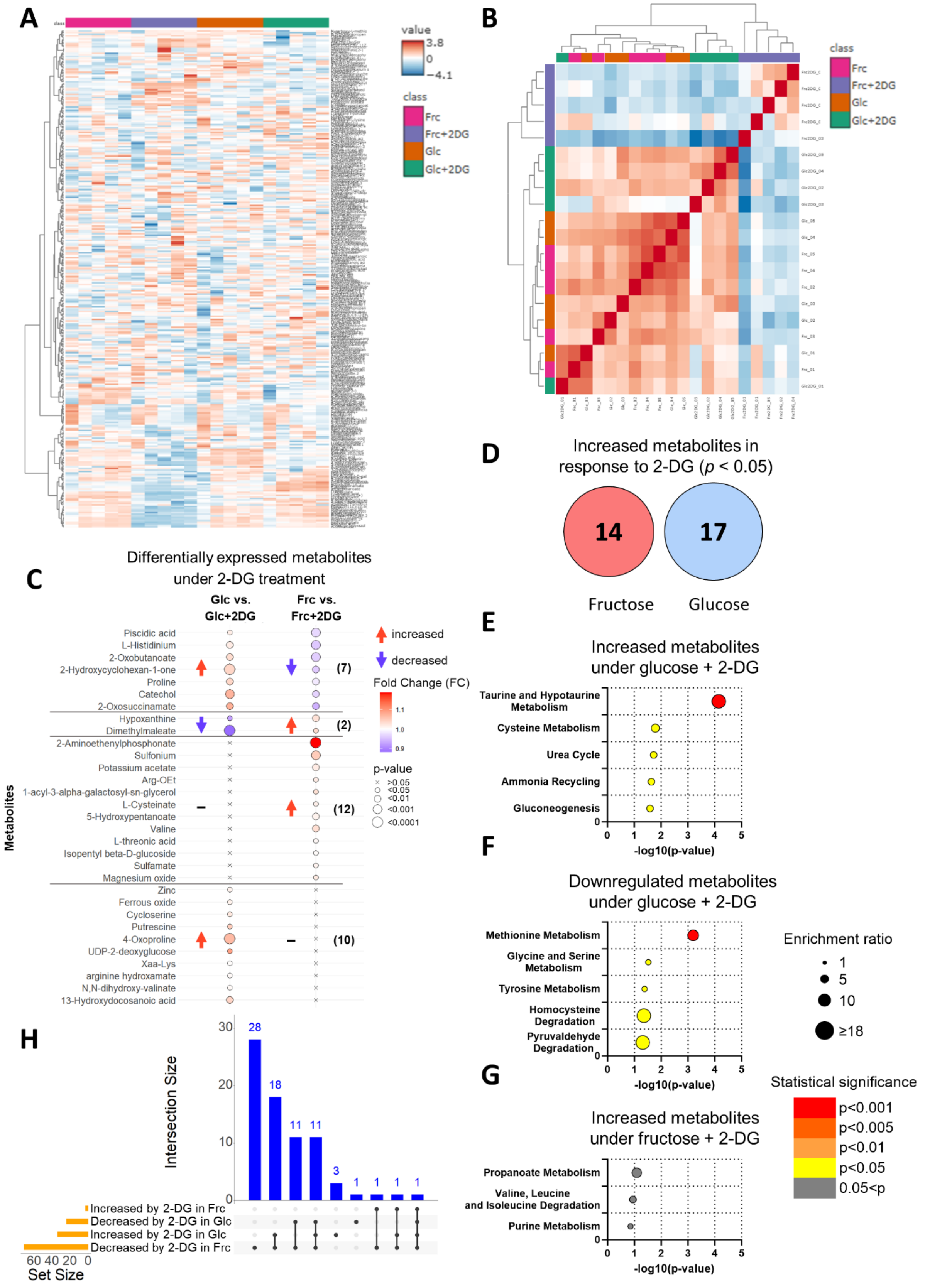

**Supplementary Figure 4. Global metabolomics revealed hexokinase independent metabolic shifts, both relevant and irrelevant of hexose condition. A.** Heatmap showing intracellular metabolomic profiles of cells cultured under four conditions:  fructose (pink), fructose + 2-DG (purple), glucose (orange), and glucose + 2-DG (green). **B.** Correlation heatmap illustrating relationships among samples from the same conditions. **C–D** Metabolomic comparisons reveal enriched metabolites with 2-DG treatment. Dot plot indicates fold-change and significance of metabolites. The numbers in parentheses indicate the number of metabolites (C). Venn diagram depicting the number of metabolites elevated following 2-DG treatment (D). **E–G.** Dot plots showing pathway enrichment of metabolites altered by 2-DG treatment under glucose-supplemented (E–F) or fructose-supplemented (G) conditions. **H.** Upset plot summarizing overlaps between differentially expressed metabolites across conditions (fructose, Frc; glucose, Glc). Collectively, the analysis reveals both hexose-dependent and -independent shifts in intracellular metabolism in response to 2-DG treatment.

#### Supplementary Figure 5

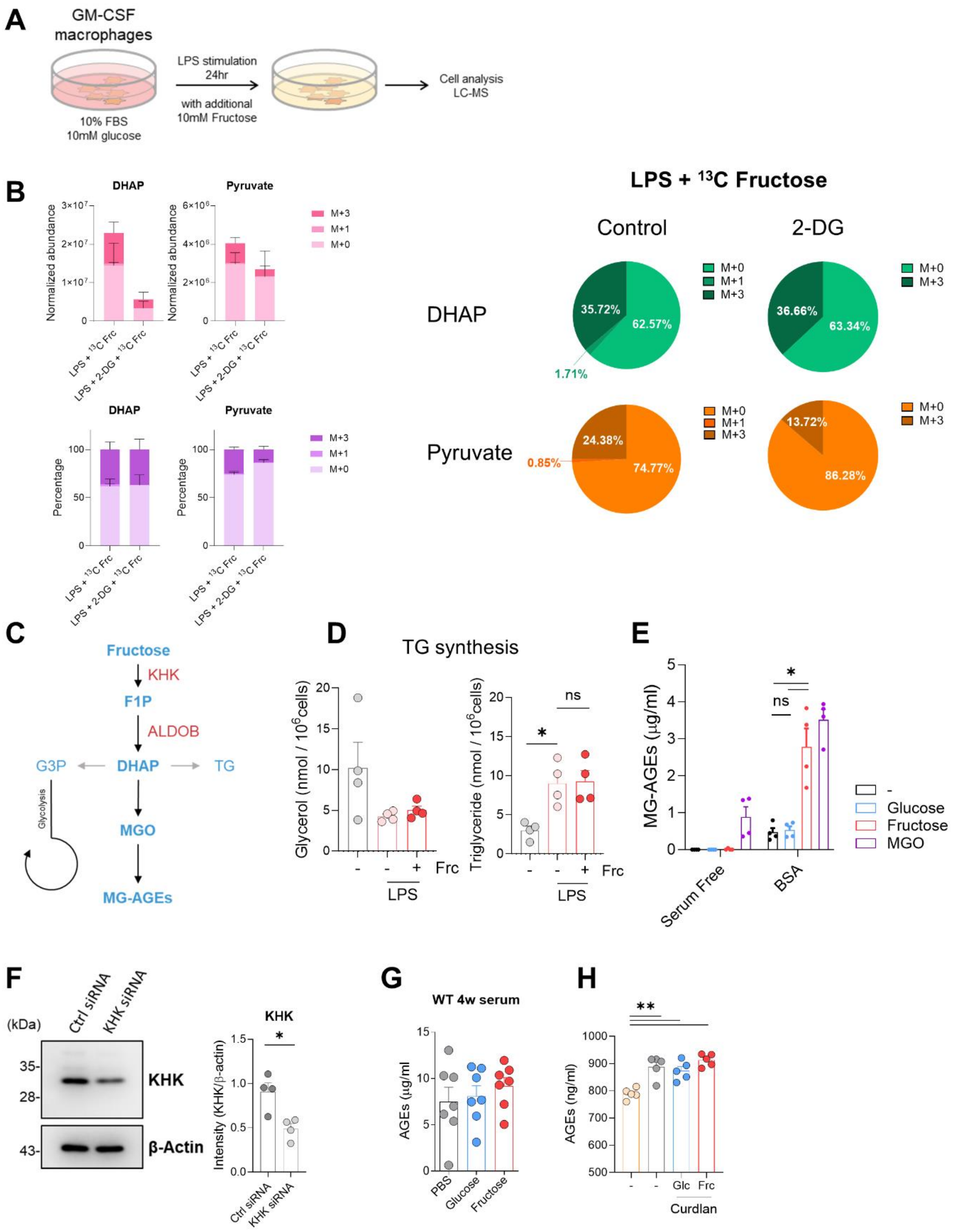

**Supplementary Figure 5. Fructose-derived dihydroxyacetone phosphate (DHAP) enhances MG-AGE formation. A.** Experimental schematic corresponding to Figure 4. **B.** Relative intensities of $^{13}C$-labeled and unlabeled DHAP and pyruvate in macrophages treated with $^{13}C_6$-fructose for 24 h in complete medium containing 50 ng/mL LPS, in the presence or absence of 0.5 mM 2-DG (n = 8). Bars represent total metabolite abundance. **C.** Overview of KHK-dependent fructose metabolism. **D.** *De novo* glycerol and triglyceride synthesis measured in cell lysates under the indicated conditions. **E.** Quantification of MG-H1 levels in culture supernatants in the presence or absence of 100 μg/mL BSA. **F.** Immunoblot analysis of KHK protein after siRNA-mediated knockdown in GM-CSF macrophages. Representative blot (left) and densitometric quantification normalized to β-actin are shown. **G.** Serum total AGE levels following 4 weeks of glucose or fructose administration in GM-CSF–treated WT mice. **H.** Serum total AGE levels in SKG mice 6 weeks after curdlan injection and hexose supplementation. Data are shown as mean ± SEM. *$p < 0.05$ and **$p < 0.01$ (Mann–Whitney *U* test).

### Supplementary Figure 6

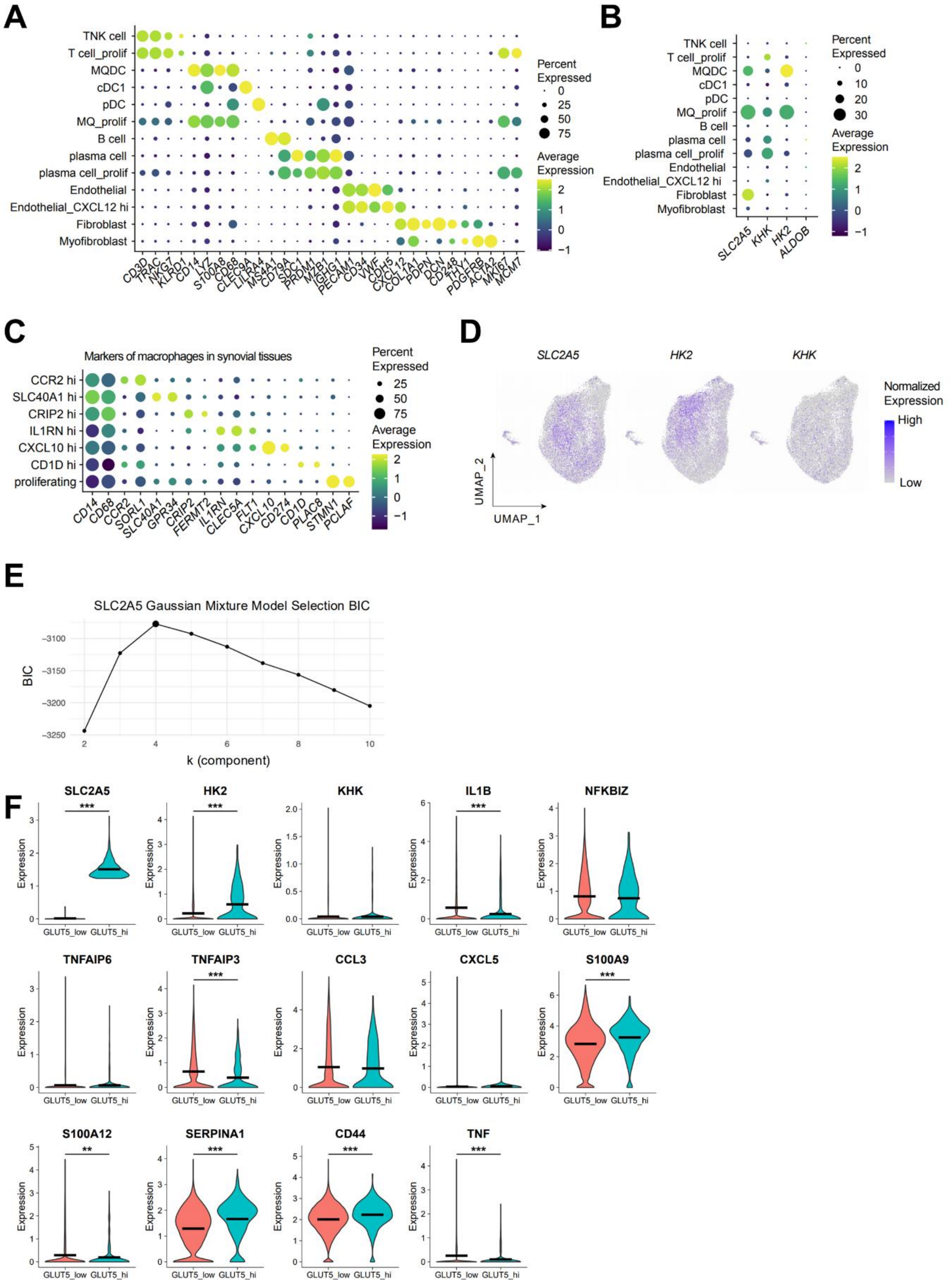


85

**Supplementary Figure 6. Fructose metabolism defines distinct transcriptomic states in RA synovial macrophages (RASFMO). A.** Dot plot showing scaled average expression of canonical marker genes across clusters identified in Figure 7C; dot color indicates expression intensity, and size denotes the proportion of expressing cells. **B.** Expression patterns of fructose metabolism–related genes across cell subsets, visualized as dot plots showing scaled mean expression and proportion of expressing cells. **C.** Dot plot of macrophage cluster marker genes corresponding to Figure 7G, illustrating transcriptional diversity among subsets. **D.** Projection of fructose metabolism–associated gene expression onto t-SNE plots (FeaturePlots). **E.** Gaussian mixture model (GMM)–based classification of macrophages according to *SLC2A5* expression; candidate cluster numbers (k) were compared using the Bayesian Information Criterion (BIC). **F.** Violin plots displaying gene expression distributions across macrophage clusters. These analyses demonstrate that fructose metabolism–associated genes delineate transcriptionally and functionally distinct macrophage populations within RA synovial tissue.

#### Supplementary Figure 7

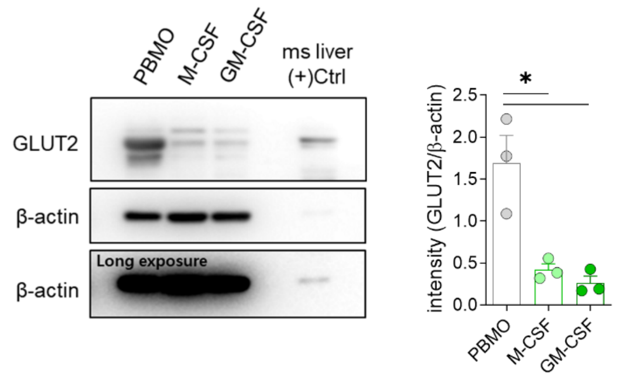


**Supplementary Figure 7. GLUT2 expression during macrophage differentiation.** GLUT2 protein expression was assessed in human monocytes from healthy controls (HCs) differentiated with M-CSF (50 ng/mL) or GM-CSF (50 ng/mL) for 6 days (n = 3). A representative immunoblot is shown (left), and the corresponding quantification is presented as mean ± SEM. *$p < 0.05$ by unpaired *t*-test.
